# Reprogramming the EnvZ-OmpR two-component system confers ethanol tolerance in *Escherichia coli* by stabilizing the outer membrane and altering iron homeostasis

**DOI:** 10.1101/2025.05.01.651661

**Authors:** Thomas Schalck, Sarah De Graeve, Lars Roba, Julia Victor Baldoma, Toon Swings, Bram Van den Bergh, Jan Michiels

**Affiliations:** Centre of Microbial and Plant Genetics, KU Leuven, Leuven, Belgium; Center for Microbiology, VIB-KU Leuven, Leuven, Belgium

**Keywords:** ethanol tolerance, *Escherichia coli*, EnvZ-OmpR two-component system, outer membrane stabilization, iron metabolism and uptake

## Abstract

Ethanol is a fermentation product widely used as a fuel and chemical precursor in various applications. However, its accumulation imposes severe stress on the microbial producer, leading to significant production losses. To address this, improving a strain’s ethanol tolerance is considered an effective strategy to enhance production. In our previous research, we conducted an adaptive evolution experiment with *Escherichia coli* growing under gradually increasing concentrations of ethanol, which gave rise to multiple hypertolerant populations. Based on the genomic mutational data, we demonstrated in this work that adaptive alleles in the EnvZ-OmpR two-component system drive the development of ethanol tolerance in *E. coli*. Specifically, when a single leucine was substituted for a proline residue within the periplasmic domain using CRISPR, the mutated EnvZ osmosensor caused a significant increase in ethanol tolerance. Through promoter fusion assays, we showed that this particular mutation stabilizes EnvZ in a kinase-dominating state, which reprograms signal transduction involving its cognate OmpR response regulator. Whole-genome proteomics analysis revealed that this altered signaling pathway predominantly maintains outer membrane stability by upregulating global porin levels and attenuating iron metabolism in the tolerant *envZ\**_L116P_ mutant. Moreover, we demonstrated that the hypertolerant *envZ\**_L116P_ allele also promotes ethanol productivity in fermentation, providing valuable insights for enhancing industrial ethanol production.

**AUTHOR SUMMARY:** Ethanol is a versatile chemical with many applications, but producing it in high quantities remains a challenge. This is because *Escherichia coli*, a candidate ethanol production strain, is naturally sensitive to this short-chain alcohol, especially when levels are gradually accumulating during fermentation. To resolve this bottleneck, we have investigated how *E. coli* can acquire tolerance to its own toxic fermentation product. Our research indicated that a single amino acid substitution in EnvZ-a key sensor protein that normally protects *E. coli* against extreme osmotic stress-is sufficient to confer ethanol tolerance. Further analysis revealed that the mutation perturbs the EnvZ-mediated signaling cascade, which, in turn, changes the transporter composition in the outer membrane and attenuates the cell’s iron metabolism. These adaptations enable *E. coli* to survive under high-ethanol conditions, thereby promoting its ethanol production efficiency. This discovery provides a suitable strategy to increase ethanol titers in industrial settings using fermentation.

## INTRODUCTION

The global climate crisis today is mainly attributed to the increased use of fossil fuels and petrol-derived products and the associated increase in high greenhouse gas (GHG) emissions [1,2]. To limit the widespread impact of global warming and reduce GHG emission-related health issues, producing biofuels from renewable resources (*e.g.*, agricultural residues or energy crops) instead of extracting crude oil from oil reservoirs is considered the most adequate strategy [3–5]. Particularly, bioethanol has become an established petroleum alternative in the US, Brazil, and the EU as transportation fuel [6]–[8] but also as a precursor for bulk chemicals such as ethylene oxide [6,7].

Although microbial-based bioethanol production is a cost-effective method, maximizing product titers often remains challenging as the microbial producer (*e.g. Saccharomyces cerevisiae*, *Zymomonas mobilis*, or *Escherichia coli* [8–11]) experiences stress from high concentrations of (lignocellulosic) biomass-derived inhibitors and end-product (*e.g.,*ethanol) [12–15]. Due to its amphiphilic character, ethanol can interact with the microbial plasma membrane, resulting in severe membrane disorders and disruption of the proton motive force and ion/nutrient fluxes [16–20]. Furthermore, this small, two-carbon alcohol may cross the membrane and, as a result, impede transcription and translation, damage protein structures and DNA (whether or not due to accumulation of reactive oxygen species) [20–25].

Previous research has indicated that adaptation to ethanol stress is a complex process in which a combination of cell envelope--adapting, ROS-scavenging, protein-refolding and, energy-restoring pathways are involved [20,23,25–29]. Here, we focus on a crucial adaptation mechanism that conserves the outer membrane (OM) structure under lethal ethanol stress. Based on the mutational dataset of a previously conducted evolution experiment under continuous ethanol stress, the amino acid substitution, L_116_P, in the central EnvZ histidine kinase osmosensor was found to confer high ethanol tolerance in *E. coli* [30,31]. We show that this particular amino acid alteration inside the periplasmic domain (PD) disturbs the native kinase/phosphatase activity. As a result, this evolved *envZ* allele reprograms the expression pattern of downstream-regulated genes and, consequently, represses the enterobactin biosynthesis and transport systems and adjusts the porin level within the outer membrane to survive lethal ethanol concentrations. Furthermore, we demonstrated that this EnvZ/OmpR-involved tolerance mechanism is also a suitable strategy to accelerate ethanol production parameters of *E. coli* in fermentation settings.

## RESULTS

### The EnvZ-OmpR two-component system underlies adaptation to lethal ethanol stress

We have previously evolved 16 *E. coli* populations (HT1-HT16) by gradually increasing the imposed ethanol concentration to obtain high-tolerant *E. coli* lines that can eventually grow in 8.5% EtOH (Figure S.1 provides an overview of the course of ethanol adaptation up to 8.5%) [30,31]. To prioritize the most relevant tolerance-conferring adaptation mechanisms in *E. coli*, we here first conducted a Gene Ontology (GO) enrichment analysis on the evolution dataset of Swings *et al.* (2017) [30]. We specifically focused our analysis on the 2,560 genes, exhibiting at least one mutation across the 16 parallel-evolved populations at each sequenced time point. This Gene Ontology (GO) enrichment analysis revealed that, throughout evolution, *E. coli* predominantly acquired mutations in genes that encode membrane-associated proteins with a role in transport or kinase-mediated signal transduction (Figure S.2). We further focused on the relevance of the *E. coli* Two-Component Systems (TCSs) in ethanol adaptation since the cellular localization and function of these bacterial signaling pathways correspond to the enriched GO terms (Figure S.2). More specifically, a bacterial TCS is composed of a membrane-associated sensor that, depending on the environmental conditions, (de)phosphorylates its cognate response regulator to induce or repress expression of downstream-regulated genes [32–34] (Figure 1B). Interestingly, mutations in the sensor and response regulator moiety of 26 distinct TCSs were frequently observed among all ethanol-evolved *E. coli* populations at some stage of adaption, especially in the EnvZ-OmpR, DpiBA, RstBA, BaeSR, and AtoSC signal transduction pathways (Figure S.3).

**Figure 1:**
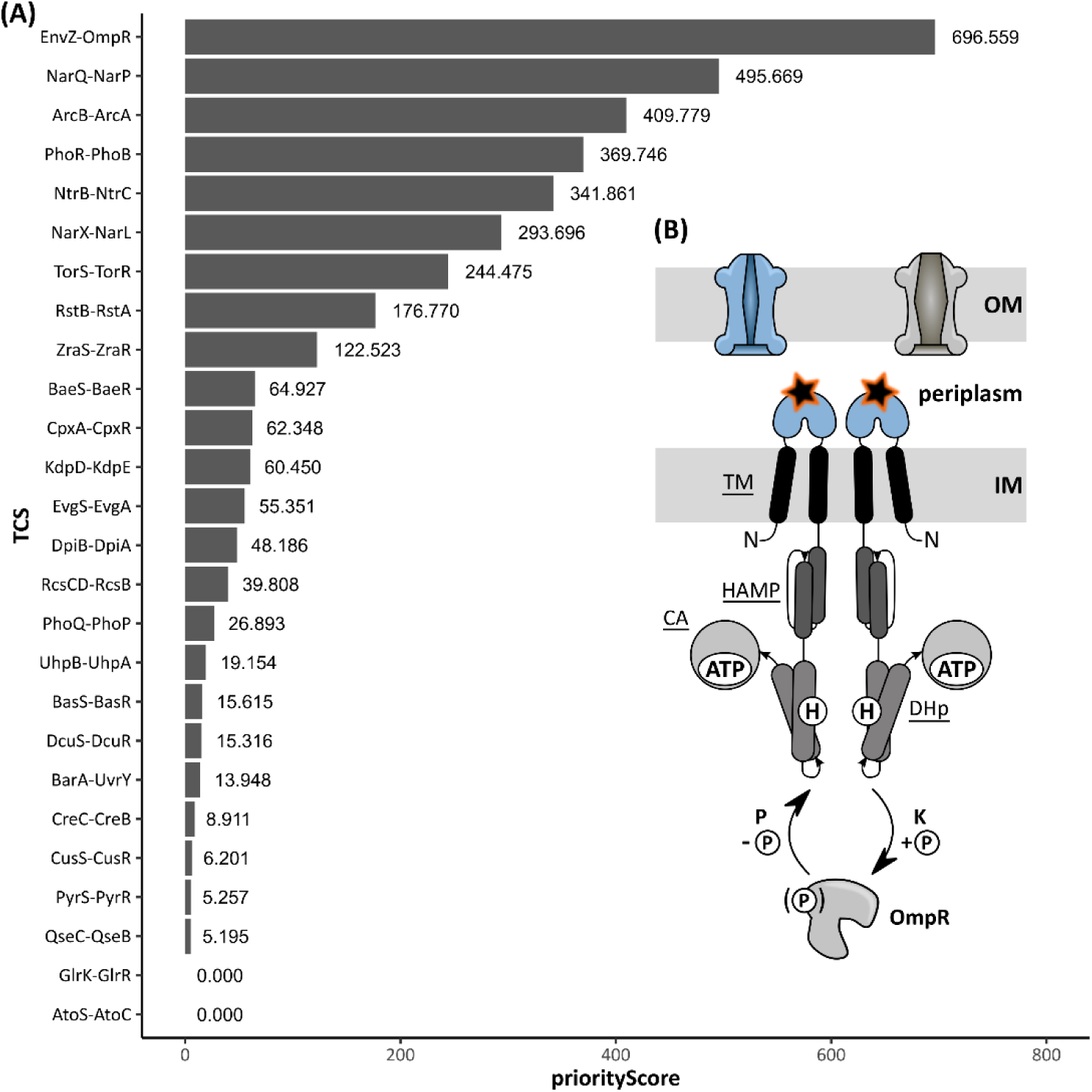
Mutation frequency and Gene-Ontology enrichment analyses characterize the EnvZ-OmpR Two-Component System as the prime evolutionary target in conferring increased tolerance towards ethanol. **(A)** The EnvZ-OmpR signal-transduction system was appointed the most relevant TCS in ethanol tolerance based on the priority score. This score was defined as the product of the mutation frequency (or coverage) across all evolved populations and the number of membrane-associated transporters within the corresponding regulon. **(B)** Schematic representation of the EnvZ-OmpR TCS. From N-to C-terminus, the EnvZ osmosensor consists of five principal domains: the transmembrane (<u>TM</u>), periplasmic, the <u>HAMP</u> (which is shared among Histidine kinases, Adenylate cyclases, Methyl-accepting proteins, and Phosphatases), the Dimeric, Helical domain containing the central H_243_ (encircled H) that is phosphorylated (<u>DHp</u>) and the C-terminal catalytic domain that binds ATP (<u>CA</u>) [33]. A hyperosmotic shock induces the kinase (K) activity of EnvZ which involves auto-*trans*-phosphorylation of H_243_ and phosphotransfer of the phosphate (encircled P) towards the D_55_ residue of OmpR [34–36]. Contrary, a hypoosmotic condition stimulates EnvZ’s phosphatase activity which removes the phosphate from OmpR [39,40]. The L_116_P amino acid substitution is depicted as a black, orange highlighted star, located at the periplasmic domain.

Based on the frequency at which mutations emerged in the sensor and response regulator moieties (Figure S.3) and the number of membrane-associated transporters within the TCS’s regulon, an overall priority score was finally assigned to each TCS (Figure 1A). The EnvZ-OmpR signaling pathway was the highest-ranked evolutionary target. Therefore, we decided to investigate the relevance of this TCS in ethanol tolerance and focused on the impact of the most frequently occurring *envZ* allele, called L_116_P. The native gene product, EnvZ, acts as a histidine kinase/phosphatase osmosensor that tunes the phosphorylation state of its cognate response regulator OmpR, according to the environmental osmolarity [32,34–38]. In turn, the degree of phosphorylation determines the DNA binding affinity of OmpR and, thus, defines the expression pattern of all downstream-regulated genes involved in the response towards extreme osmotic stresses [39,40]. Given the emergence of the L_116_P substitution in different parallel populations and the pivotal role of EnvZ in membrane protein regulation, we anticipated that the mutant EnvZ*_L116P_ osmosensor may confer ethanol tolerance by remodeling the cell envelope structure as a result of altered regulation of membrane-associated (transporter) proteins.

### The L_116_P mutation in *envZ* is adaptive and improves ethanol tolerance *in vivo*

To verify whether the L_116_P allele of interest is sufficient to improve ethanol tolerance, the mutation was reconstructed in the wild-type background. The growth kinetics and cell survival of *envZ\**_L116P_ were subsequently assessed in the presence of 5% ethanol. The latter corresponds to the concentration at which cell growth of the wild-type BW25113 strain is completely inhibited and at which the evolution experiment was originally initiated [30]. Additionally, strains lacking either *envZ* or *ompR* were also subjected to 5% ethanol stress (Figure 2). Strikingly, the *envZ\**_L116P_ mutant is able to grow in a 5% ethanol-enriched medium and survives an otherwise lethal ethanol stress. In contrast, disrupting EnvZ and OmpR results in a strong decrease in both microbial growth and survival (Figure 2). Hence, OmpR mediates the tolerance-conferring property of the mutant EnvZ*_L116P_ osmosensor since deleting this response regulator completely abolishes the growth and survival advantage of the gain-of-function L_116_P allele.

**Figure 2:**
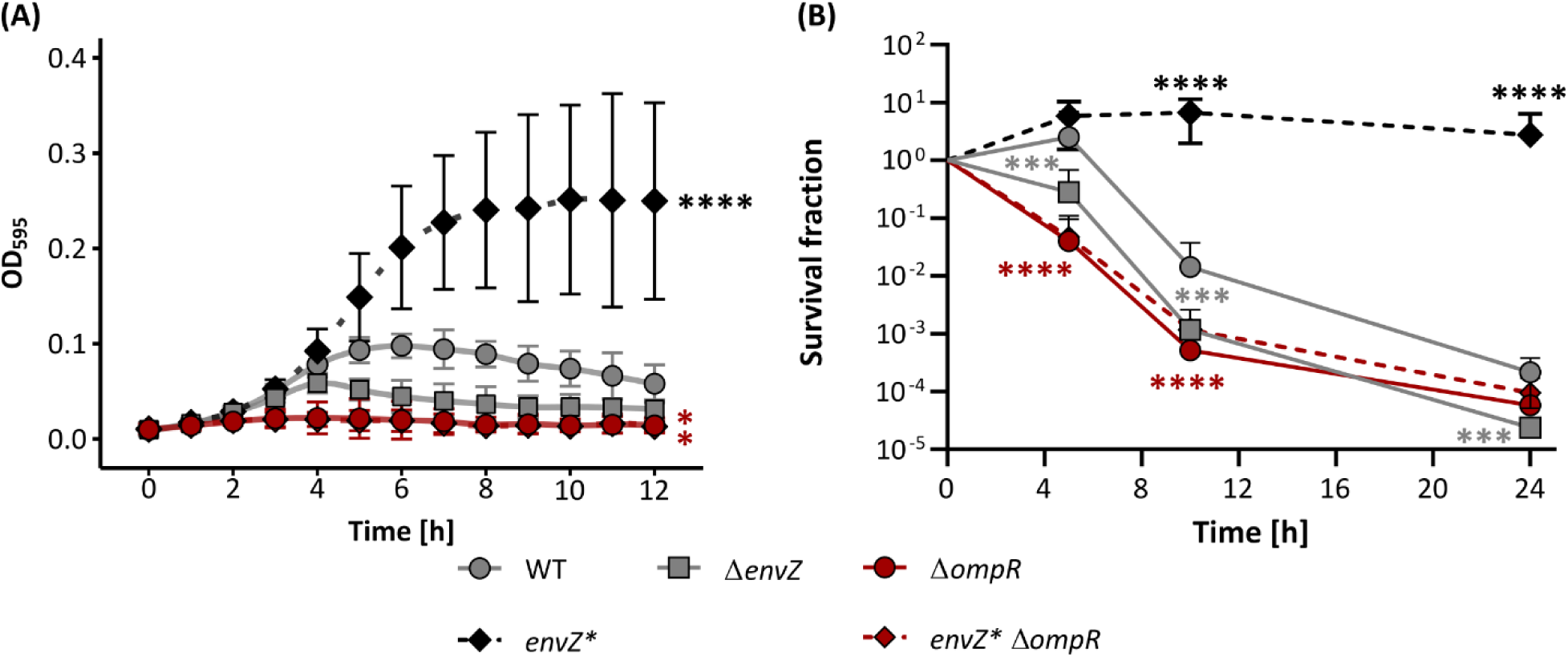
The L_116_P allele improves ethanol tolerance in an OmpR-dependent way. **(A)** The splines-fitted growth curves for the BW25113 wild type (WT), the *envZ\**_L116P_ mutant, and EnvZ-OmpR deletion mutants (Δ*envZ* and Δ*ompR*) under a 12h exposure to 5 (v/v)% ethanol. Based on the QurvE-fitted growth curves [41], the optical density (OD_595_) at the plateau phase and starting point (t_0_) were extracted (Figure S.5). The OD_595_ increment at the plateau phase *vs.* the start (t_0_) for each strain was considered to calculate the statistics. *P*-values were obtained from one-way ANOVA with a Dunnett’s posthoc test. Error bars represent the 95% confidence intervals (n=4). **(B)** Survival fraction of the corresponding *E. coli* strains. Data points indicate the mean and error bars represent the 95% confidence intervals (n=4). Levels of significance are indicated as follows: *, *P*≤0.05, **, *P*≤0.01, ***, *P*≤0.001, ****, *P*≤0.0001 and are derived from a Generalized Linear Interaction Model with Time [h] as a (discrete) factor using the WT strain as the baseline, paired with a Dunnett’s post-hoc test. The *P*-values of the Δ*ompR* strains are identical and highlighted in red.

### Kinase-promoting *envZ* variants confer ethanol tolerance

Given the central role of EnvZ in osmoregulation and -tolerance [42], we anticipated that the L_116_P substitution resulted in a reprogrammed bifunctional kinase:phosphatase activity of EnvZ and, therefore, caused the improved tolerance phenotype. To compare the kinase:phosphatase activity of the mutant with the wild-type EnvZ sensor, we recorded the expression pattern of *ompC* and *ompF*, encoding the major outer membrane porins, using a fluorescence-coupled promoter fusion assay [43–45]. Previous research has indicated that *ompC* is induced or repressed and *ompF* repressed or induced when OmpR-P levels are high or low, respectively (Figure 3C) [39].

**Figure 3:**
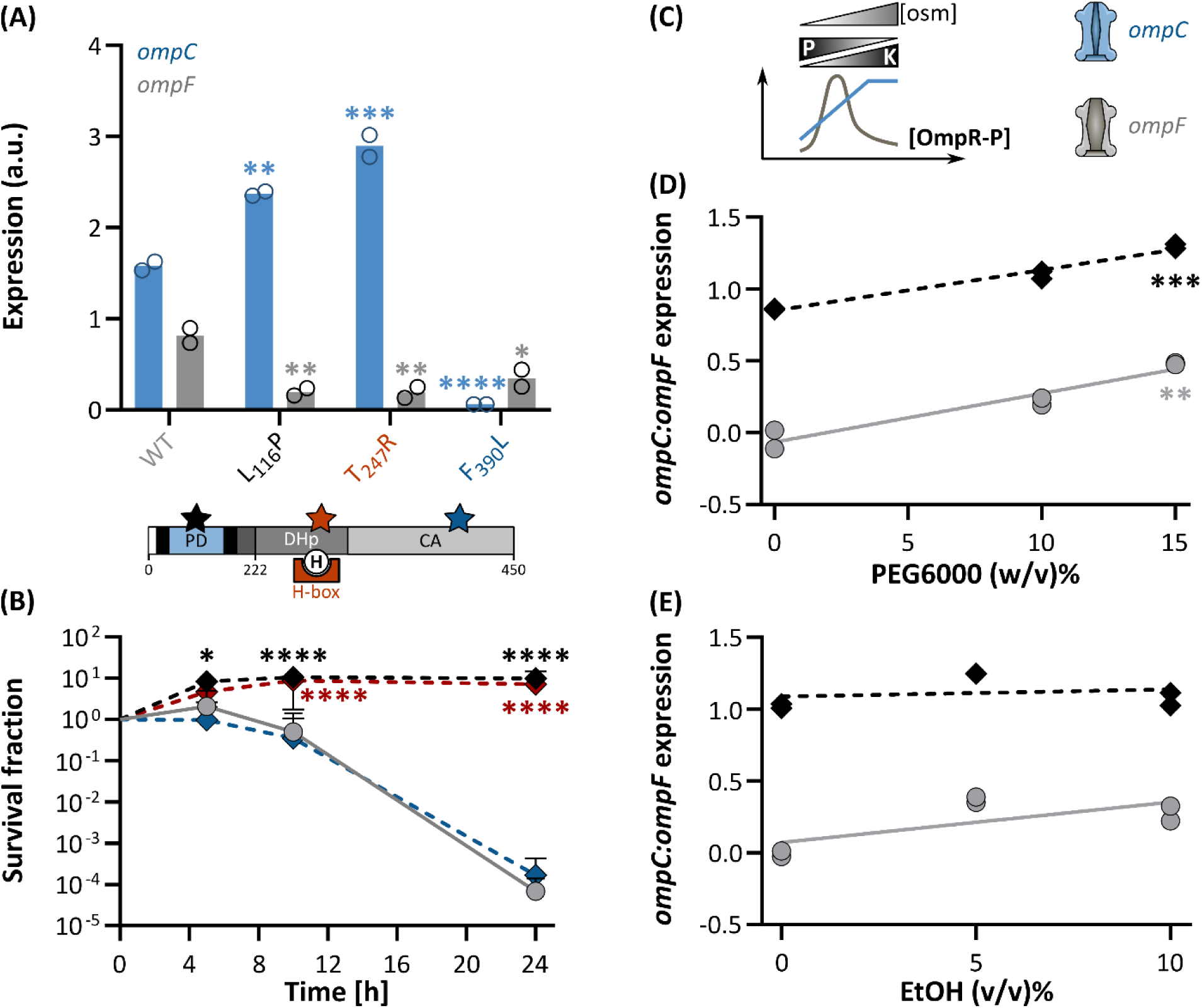
Constitutive, kinase-promoting *envZ* substitutions improve tolerance in *E. coli.* **(A)** (Top) When the kinase-promoting L_116_P and T_247_R are present in the EnvZ osmosensor, *ompC* (blue) is upregulated and *ompF* (grey) is repressed, compared to the wild type. In contrast, the kinase-defective EnvZ*_F390L_ induces neither *ompC* nor *ompF*. (Bottom) Schematic representation of the EnvZ domain structure (PD, periplasmic domain; H-box, the amino acid sequences surrounding the central H_243_; EnvZc, catalytic C-terminal moiety encompassing the DHp and CA domains, the other references match the nomenclature in Figure 1) together with the ethanol-evolved L_116_P mutation (black star) and literature-derived T_247_R (red star) and F_390_L (blue star) substitutions [46–48]. Levels of significance are derived from a (Generalized) Linear Mixed Effects Regression model with Dunnett’s post-hoc test (with WT as the reference). **(B)** The impact of reprogramming the enzyme activities of EnvZ on ethanol tolerance, expressed in terms of survival. The *envZ*\*_L116P_ and *envZ*\*_T247R_ mutants that show elevated *ompC* levels (*i.e.* display high kinase:phosphatase ratios), can cope with ethanol stress significantly better than *envZ*\*_F390L_ or *envZ*_WT_ that do not share the increased *ompC* expression feature. Levels of significance are derived from a Generalized Linear Mixed Effects Regression model followed by a Dunnett’s post-hoc test (with WT as reference). **(C)** The anticipated OmpR-P levels and OM porin expression profile in *E. coli* as a result of changing EnvZ kinase:phosphatase activities in response to the environmental osmolarity. The large pore size OmpF porin predominates at low osmolarity, corresponding to decreased kinase:phosphatase ratios and hence low OmpR-P levels. High osmolarity, in contrast, induces high kinase:phosphatase ratios, increases the OmpR-P concentration, and favors *ompC* expression, encoding an OM porin with a small diffusion pore [49]. The log_10_-transformed *ompC:ompF* expression ratio, reflecting the EnvZ kinase:phosphatase bifunctional activity, in response to hyperosmotic **(D)**, mimicked using high PEG6000 concentrations, and ethanol **(E)** stress. Expression intensities of each porin in each strain (wild type, grey and *envZ*\*_L116P_, black) were expressed in terms of FITC fluorescence values as measured through flow cytometry. Levels of significance are derived from a t-test on the slope of a Linear Regression Model. Levels of significance:*, *P*≤0.05, **, *P*≤0.01, ***, *P*≤0.001, ****, *P*≤0.0001. Raw flow cytometry histograms are provided in Figure S.8.

By tracking *ompC* and *ompF* expression levels, we demonstrated that the *envZ*\*_L116P_ mutant, in stress-free medium, stimulates *ompC* and strongly represses *ompF* with respect to the wild type (Figure 3A, Figure S.6). Hence, the EnvZ*_L116P_ osmosensor has an altered kinase:phosphatase balance in favor of the kinase activity that is reminiscent of the hyperosmotic, kinase-dominant state. Furthermore, mutants such as Δ*envZ* and Δ*ompR* that are hypersensitive towards ethanol and are dysfunctional in EnvZ-OmpR signal transduction, express both *ompC* and *ompF* barely (Figure **2**S.6C). Altogether, our findings suggest that a high kinase:phosphatase ratio and the resulting, reprogrammed signal transduction pathway underlies ethanol tolerance. To further corroborate this, we studied two *envZ* alleles that have been examined before in terms of their impact on the kinase:phosphatase equilibrium but have not been evaluated yet with regard to ethanol tolerance (Figure 3A). The first one, T_247_R, is located in the DHp-associated H box, nearby the catalytic H_243_, and is known to favor EnvZ’s kinase state, like the L_116_P allele [46,47]. In contrast, the CA domain-located F_390_L substitution renders EnvZ kinase-deficient and simultaneously increases its phosphatase activity [48]. While the *envZ*\*_T247R_ mutant, similar to *envZ*\*_L116P_, displays improved tolerance, the *envZ*\*_F390L_ strain is equally hypersensitive to ethanol as compared to the ancestral wild type (Figure 3B).

Next, we subjected the ethanol-tolerant *envZ*\*_L116P_ mutant to ethanol and hyperosmolarity to assess whether the evolved EnvZ*_L116P_ osmosensor is ‘locked’ in a constitutive high kinase state irrespectively of the environmental stressor and, thus, whether the ethanol-adapted system is still responsive to stresses. To impose a hyperosmotic stress, we mixed polyethylene glycol (PEG6000) as osmolyte into the growth medium to, ultimately, force EnvZ into its high kinase:low phosphatase state (Figure 3C, Figure S.7). Consequently, the latter should be reflected in a high *ompC*:*ompF* expression ratio. In response to hyperosmotic stress, the L_116_P allele displays native osmosensing ability as *ompC* expression is triggered slightly and *ompF* expression is repressed strongly in accordance with the PEG6000 concentration, similar to the wild-type EnvZ (Figure S.8, Figure 3D). As a result, the *ompC*:*ompF* ratio, albeit consistently higher in the *envZ*\*_L116P_ mutant, increases as would be expected in response to hyperosmotic stress (Figure 3D). In contrast, ethanol stress did not evoke any change in biochemical activity neither in the wild type nor in the ethanol-evolved osmosensor (Figure 3E, Figure S.8). The latter suggests that the EnvZ osmosensor, regardless of the L_116_P substitution, is inherently insensitive to ethanol. Therefore, cells can only benefit from improved ethanol tolerance when they acquire the L_116_P allele that forces the EnvZ osmosensor in a constitutive, ethanol-independent K>P state.

### The outer membrane porin, OmpC contributes to ethanol tolerance in the *envZ*\*_L116P_ mutant

The EnvZ-OmpR-tandem acts as a signal transduction cascade in which the OmpR response regulator ultimately adjusts the expression pattern of multiple genes that belong to the OmpR regulon, according to the kinase:phosphatase balance of EnvZ [50]. Importantly, eliminating *ompR* turns both the wild-type and *envZ*\*_L116P_ mutant strains hypersensitive to ethanol, indicating that at least one OmpR downstream-regulated gene should be involved in the ethanol tolerance phenotype. Therefore, we first evaluated the role of OmpC and OmpF in ethanol tolerance since the expression of both porins was significantly altered in the *envZ*\*_L116P_ mutant (Figure 3A). In general, inactivating *ompC* drastically compromised growth (Figure 4A, B) and survival (Figure 4D, E) under ethanol stress, although the effect was less pronounced compared to the inactivation of *ompR* (Figure 4C). However, deleting *ompC* did not fully abolish the *envZ*\*_L116P_-associated tolerance phenotype since the survival of *envZ*\*_L116P_ Δ*ompC* mutant was still higher than the WT reference strain with *ompC* after 24h of ethanol exposure (*P*<0.0001). In contrast to eliminating *ompC*, removing *ompF* has no impact on ethanol tolerance (Figure 4 A, B). Alongside the survival-based tolerance assay, we reached the same conclusions on ethanol tolerance or sensitivity when the cell shape was microscopically monitored over time in a differentially-labeled, coculturing ethanol exposure experiment (Figure S.11). While ethanol-induced morphological deformations were limited in *envZ*\*_L116P_ and *envZ*\*_L116P_ Δ*ompF* (Figure S.11A, B, C, I, J, and K), the wild-type, Δ*ompC*, and *envZ\**_L116P_ Δ*ompC* strains exhibited elongated cell shapes (Figure S.11 E, F, and G), reflecting their hypersensitivity for ethanol stress. Moreover, in the Δ*ompC* strains, subpopulations emerged that were unable to stably express their fluorescent label when exposed to prolonged ethanol stress, suggesting a deficit in gene expression (Figure S.12). Hence, OmpC is an important member of the OmpR regulon involved in ethanol tolerance.

**Figure 4:**
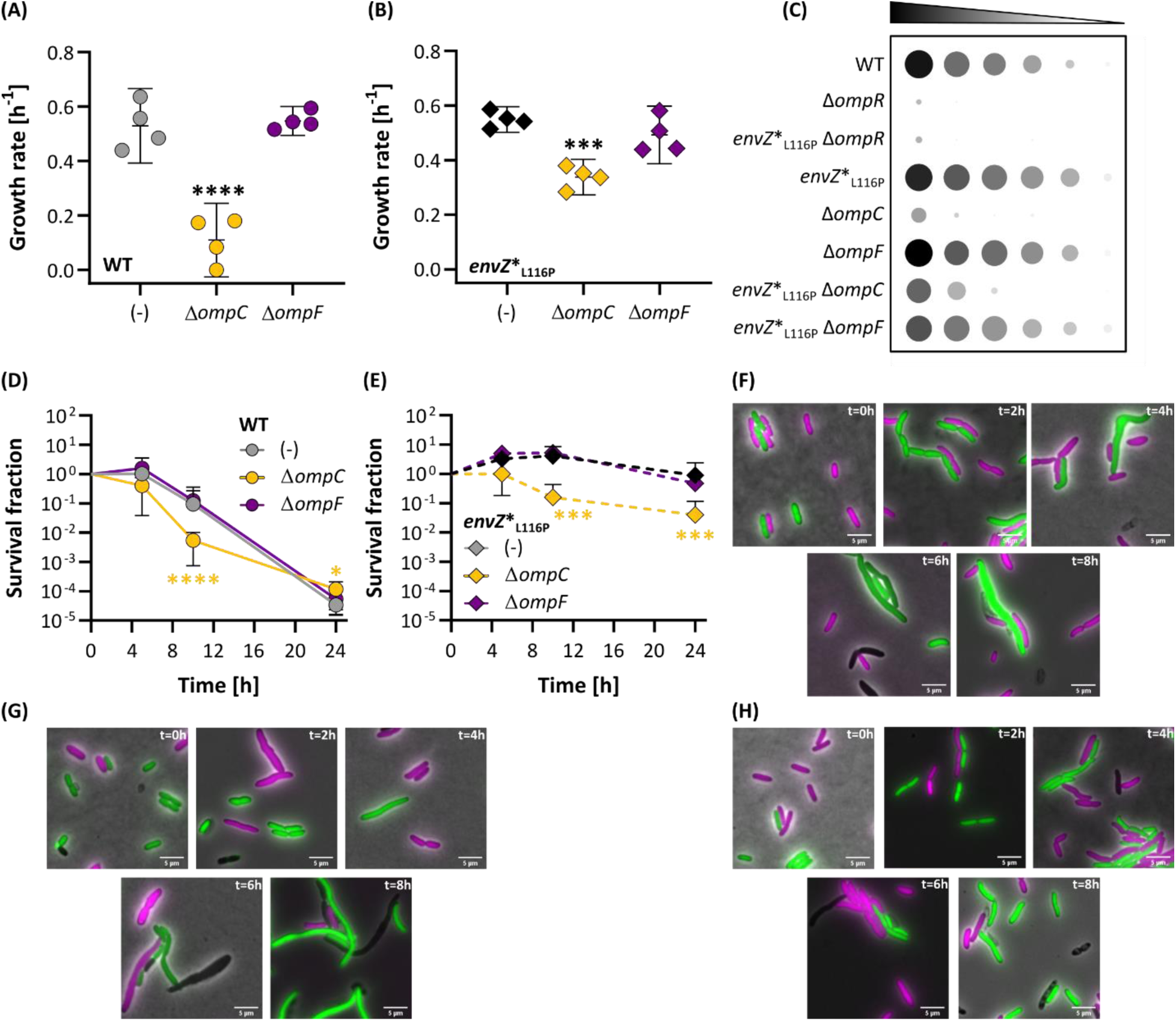
Deleting *ompC* affects survival and cell morphology under ethanol stress, whereas removing *ompF* does not compromise these parameters. The growth rate of the wild-type ancestor **(A)** or *envZ*\*_L116P_ **(B)** lacking either the *ompC* or *ompF* porin genes. (−) indicates either the wild type or tolerant mutant with intact porins. Growth rates were determined from OD-curves (at 595 nm) as shown in Figure S.9 and statistically compared using a one-way ANOVA with Dunnett’s post-hoc test (in which either the WT, A, or *envZ*\*_L116P_, B, serve as a reference). **(C)** Quantitative summary representing the mean spot area and whiteness intensity of four independent assays including eight different strains. Bacteria were diluted up to 10^6^-fold and spotted on LB agar containing 6% ethanol. The results of the 10^6^ dilution were excluded from this picture since barely any bacterial growth could be detected. Each bacterial spot was quantitatively analyzed in ImageJ (see Materials & Methods). The original images are displayed in Figure S.10. The effect of deleting *ompC* or *ompF* on the survival under ethanol stress, measured by CFU enumeration, of the wild-type **(D)** or the *envZ*\*_L116P_ strain **(E)**. Statistics of (D) and (E) are derived from a Generalized Linear and Linear Model (based on the AIC outcome), respectively, paired with a Dunnett’s post-hoc test. **(F-H)** Microscopy image samples of a mixed WT (green) - *envZ*\* (magenta) **(F)**, *envZ*\*_L116P_ Δ*ompC* (green) – Δ*ompC* (maganta), and (H) *envZ*\*_L116P_ Δ*ompF* (green) – Δ*envZ*\*_L116P_ (magenta) populations, exposed to 5% ethanol for 8h. The white line at the bottom right of each image represents the 5 µm reference scale bar. A detailed analysis of the microscopy images can be found in Figure S. 11.

### OmpF rescues the ethanol sensitivity of an ompC deletion by reducing the ethanol-induced permeabilization

Our results show that *E. coli* cells lacking OmpC exhibit a lower survival (Figure 4) and suffer from severe morphological defects under ethanol stress (Figure S.11). In contrast, OmpF was not identified as a crucial factor for improved survival under ethanol stress neither in the wild type nor in the *envZ*\*_L116P_ mutant, despite the fact that this porin shares high sequence similarity with OmpC and is also implicated in the osmotic stress response [51]. Compared to OmpC, the diffusion pore of OmpF is flanked by a lower number of charged residues increasing its pore size [44,51]. Therefore, we reasoned that the tolerance difference between the Δ*ompC* and Δ*ompF* deletion strains may be attributed either to distinctive, inherent properties of OmpC and OmpF (*e.g.* pore size) or because their absence causes the global outer membrane porin (OMP) content to change. The latter is more relevant in case of OmpC since we already provided evidence that the expression of *ompC* is at least equally high than that of *ompF* in absence of stress and even higher when the host is exposed to ethanol or hyperosmotic stress (Figure 3A, D, and E). If the global porin level in the OM is indeed critical, compensating for the lack of *ompC* in Δ*ompC* deletion mutants by providing an additional copy of *ompF* should rescue the ethanol strains under ethanol stress. Therefore, we replaced the original *ompC* gene in the wild-type and *envZ*\*_L116P_ strains with an identical, second *ompF* gene copy (called *ompF’*) to create two new strains, designated *ompF*’/*ompF* and *envZ*\*_L116P_ *ompF*’/*ompF* respectively. Importantly, the original *ompC* promoter and the 5’ UTR were preserved ensuring that *ompF’* was regulated in the same manner as the *ompC* gene it replaced. Indeed, LC-MS/MS-based quantification of the porin abundances between the *envZ*\*_L116P_ and *envZ*\*_L116P_ *ompF*’/*ompF* mutants revealed that the global porin level was identical. However, while the predominant porin in *envZ*\*_L116P_ was clearly OmpC (with very few OmpF), the outer membrane in the *envZ*\*_L116P_ *ompF*’/*ompF* mutant solely consisted of OmpF (Figure 5A).

**Figure 5:**
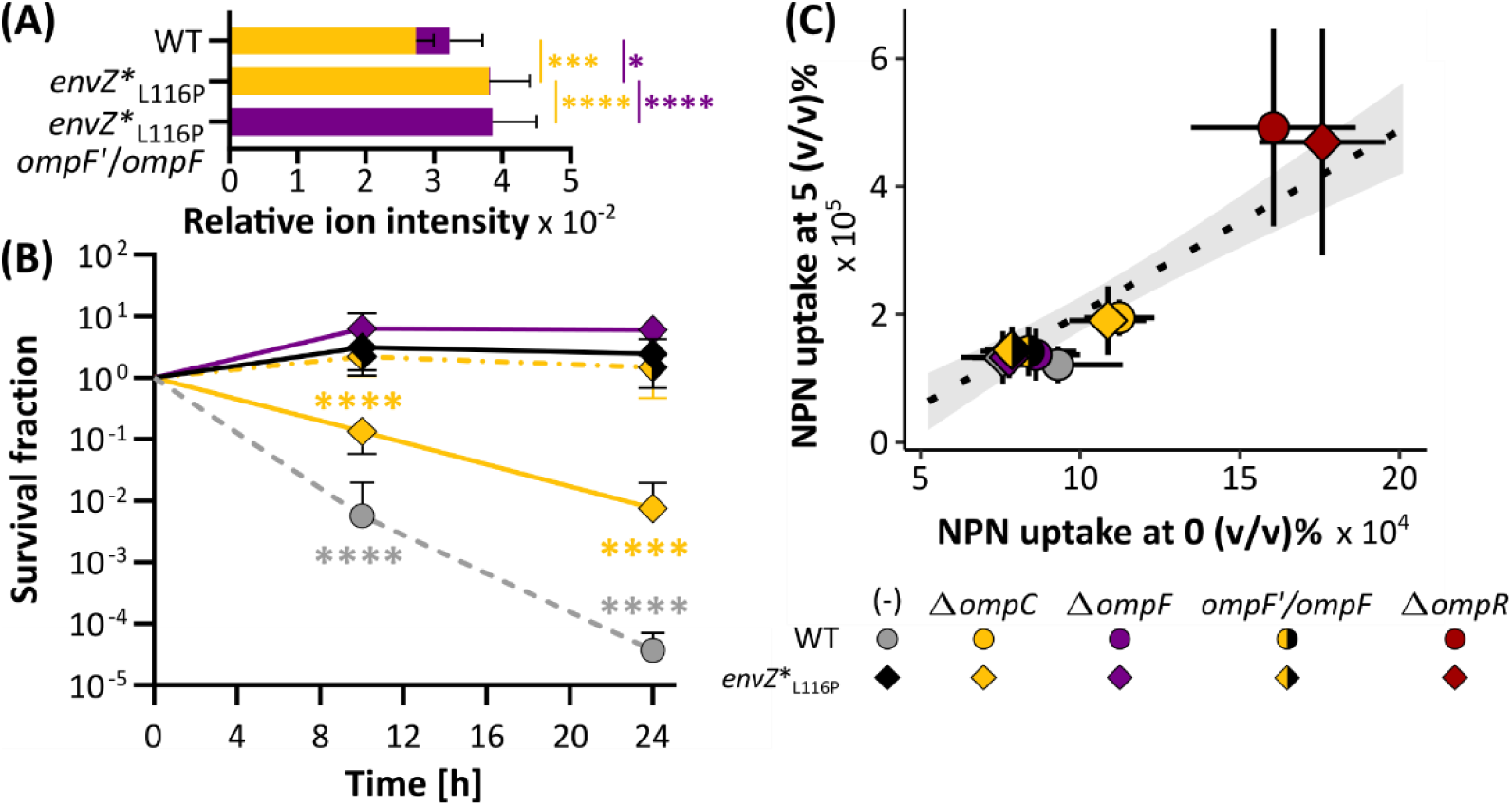
The deleterious effect of deleting *ompC* in *envZ*\*_L116P_ is reverted by integrating an extra *ompF* gene copy (*ompF’*) at the original *ompC* locus. **(A)** The relative ion intensities (or counts), defined as the normalized label-free quantification (LFQ) values of each porin (for each replicate), as a proxy for relative porin abundance. Yellow represents OmpC whereas purple indicates the OmpF level. Statistical interference is derived from one-way ANOVA combined with a Tukey multiple comparisons post-hoc test. **(B)** The *envZ*\*_L116P_ with the additional *ompF’* copy is equally tolerant as the *envZ*\*_L116P_ mutant, as opposed to *envZ*\*_L116P_ Δ*ompC.* Levels of significance are obtained from a Generalized Linear Model paired with a Dunnett’s post-hoc test using *envZ*\*_L116P_ as the reference (n=4). Error bars represent the 95% confidence intervals. **(C)** Correlation between the *N*-Phenylnaphthalen-1-amine (NPN) uptake, which is a measure for outer membrane permeability, at the 5 vs. 0 (v/v)% ethanol condition. The points represent the mean values while the crosshairs indicate the 95% confidence intervals (in x-, and y-direction). The dotted line in the background denotes a Linear Regression Model together with the grey ribbon reflecting the 95% confidence interval. Levels of significance: *, *P*≤0.05, **, *P*≤0.01, ***, *P*≤0.001, ****, *P*≤0.0001.

Moreover, the total OmpC and/or OmpF porin level in strains carrying the EnvZ*_L116P_ mutant sensor was also higher compared to the wild type (a one-way ANOVA with Tukey’s post-hoc test, *P*=0.04), indicating that its superior tolerance is linked to an overall OM content. When *ompC* was replaced with an *ompF* copy, the survival of the *envZ*\*_L116P_ mutant under 5% ethanol stress was not compromised, while just knocking out *ompC* severely diminished ethanol tolerance (Figure 5B; Figure S.13). Since outer membrane proteins, such as the β-barrel outer membrane proins OmpC and OmpF (OMPs), contribute to the structural integrity of the outer membrane in Gram-negatives, we anticipate that altering their composition has a profound impact on the cell envelope physiology and properties [52–54]. Because ethanol is known to cause cell envelope defects, reflected in the severe cell shape deformations observed in Figure S.11, we argued that a cell’s outer membrane (OM) permeability and stability, as dictated by its OMP abundance, defines its tolerance or susceptibility to ethanol stress. To corroborate this, we monitored the uptake of *N*-phenylnaphthalen-1-amine (NPN) as an indicator for ethanol-induced disruption of the OM. This dye exclusively turns fluorescent when it crosses a permeabilized OM and binds to the inner, cytoplasmic membrane [55,56]. As a proof-of-concept, the fluorescence signal intensity of the NPN dye, representing OM permeability, is dependent on the applied concentration of ethanol or polymyxyin B, a well-known antibiotic that causes OM damage (Figure S.14). This assay revealed that the OM permeability under 5% ethanol exposure globally correlates approximately with a strain’s intrinsic OM permeability (without any ethanol) (Figure 6C). Importantly, OM permeability was directly linked to a strain’s genotype. Most noticeably, deleting *ompR* severely impairs OM permeability and the same is also valid for *ompC*, although the effect is less pronounced. Providing an additional *ompF* gene copy restores the OM permeability (*i.e.* integrity) of the Δ*ompC* strains back to the baseline OM permeability of WT and *envZ*\*_L116P_, respectively (Figure 5C).

**Figure 6:**
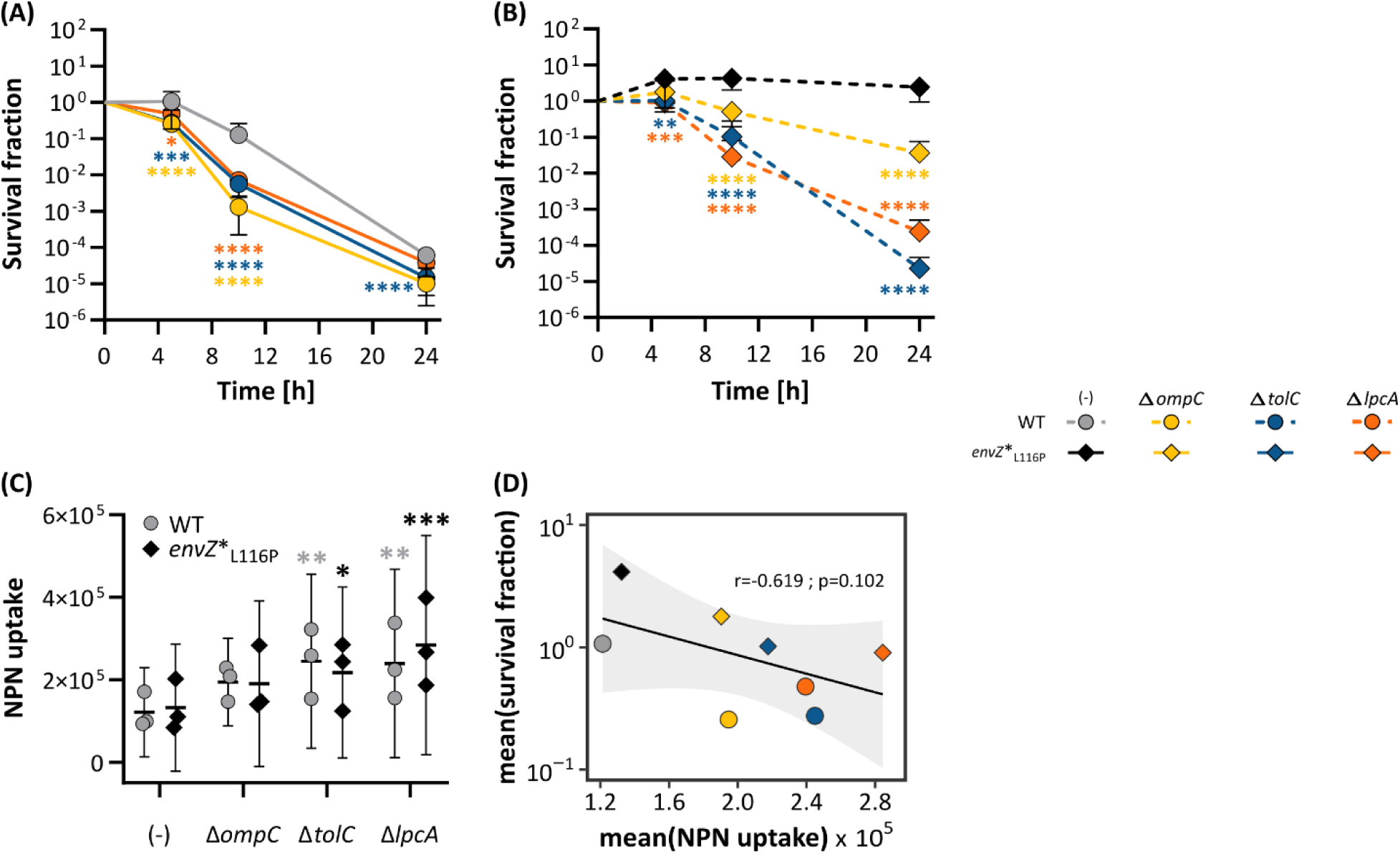
Inactivation of *tolC*, *lpcA*, and *ompC* reduces cell viability of the *envZ*\*_L116P_ tolerant mutant under ethanol stress as a result of an increase in OM permeability. Effect of deleting *ompC*, *tolC*, and *lpcA* on ethanol tolerance in the wild-type **(A)** and *envZ*\*_L116P_ mutant **(B)** strains. Statistics are inferred from a Generalized Linear Model with a Dunnett’s post-hoc test (n=4, error bars represent 95% confidence interval) using either the WT or *envZ*\* as the reference. Levels of significance: *, P≤0.05, **, P≤0.01, ***, P≤0.001, ****, P≤0.0001. **(C)** The impact of deleting *ompC*, *tolC*, and *lpcA* on the OM permeability under 5% ethanol stress (measured as NPN uptake). Statistics are derived from a Mixed Effects Linear model linked with a Dunnett’s post-hoc test with either the WT or *envZ*\*L116P as reference. **(D)** Tolerance, expressed as mean survival fraction, *vs.* the mean NPN uptake under 5% ethanol exposure for 5h (n=3). The correlation parameters are derived from a Spearman correlation test. The grey infill represents the 95% confidence interval of the black correlation curve.

In conclusion, rather than composition, outer membrane protein abundance determines ethanol tolerance, as insufficient outer membrane porins affect OM properties, like permeability, making the host more ethanol-sensitive.

### LPS biosynthesis and the TolC OM channel are implicated in EnvZ*_L116P_-mediated ethanol tolerance

We noticed that *E. coli* strains lacking OmpR are more vulnerable to ethanol stress than those with a single *ompC* deletion (Figures 2, and 4). To identify these additional tolerance-conferring genes within the OmpR regulon, we pursued two parallel approaches. First, we constructed a collection of deletion mutants in the *envZ\**_L116P_ background, each lacking a single known target of the OmpR regulon (Table S.1), and assessed the ethanol tolerance of each mutant.

The first approach identified two gene deletions that significantly impaired the ethanol tolerance phenotype of *envZ*\*_L116P_. Deletion of *lpcA*, which encodes the sedoheptulose 7-phosphate isomerase lipopolysaccharide (LPS)-biosynthesis enzyme, or *tolC*, a component of the AcrAB-TolC efflux pump-associated outer membrane channel, turn the *envZ*\*_L116P_ mutant susceptible to ethanol stress (Figure 6A, B; Figure S.15). Remarkably, eliminating the AcrA and AcrB subunits of the AcrAB-TolC tripartite extrusion system or their corresponding repressor, AcrR, does not affect the superior ethanol tolerance phenotype of *envZ*\*_L116P_ (Figure 6A, B; Figure S.15).

Since all identified OmpR hits are either an intrinsic component of the OM (*tolC*, and *ompC*) or are involved in LPS biogenesis (*lpcA*), we also studied the impact of deleting these genes on OM permeability. We observed that permeability was significantly affected in both the wild type and *envZ*\*_L116P_ mutant when *ompC,* and particularly *lpcA* or *tolC*, were deleted (Figure 6C). In addition, strains with a higher OM permeability also tend to score lower in terms of ethanol tolerance (Figure 6D).

### Reduced enterobactin biosynthesis and iron transport underly the superior tolerance of the *envZ*\*_L116P_ mutant

Aside from investigating the importance of OmpR regulon members for the ethanol tolerance phenotype, we expanded our scope to identify additional tolerance-conferring candidates using whole-genome shotgun proteomics. Gene Ontology and KEGG pathway enrichment analysis of the proteomes revealed that proteins related to iron and siderophore transport (FhuA, Fiu, FecABE, and OmpF), and enterobactin biosynthesis (EntACEF) are underrepresented in the tolerant *envZ*\*_L116P_ and *envZ*\*_L116P_ compared to the wild-type strain (Figure S.16).

To confirm the causality of the iron metabolism-involved gene set for the EnvZ*_L116P_-associated tolerance phenotype, we tested the survival of the corresponding deletion mutants under 5% ethanol stress. We hypothesized that knocking out these specific genes would bring the tolerance level closer to, or even match, the one of *envZ*\*_L116P_ since siderophore biosynthesis and iron transport gene clusters are repressed in this mutant. The subsequent survival assay (Figure 7C) revealed that deleting almost all genes within the enterobactin biosynthesis cluster, except for *entC*, along with the OM enterobactin transporter (*fepA*) enhances ethanol tolerance of the sensitive wild-type strain. In addition, knocking out *fecB*, encoding the periplasmic substrate-binding component of the iron dicitrate ABC transporter, also proved beneficial for ethanol tolerance (Figure 7C).

**Figure 7:**
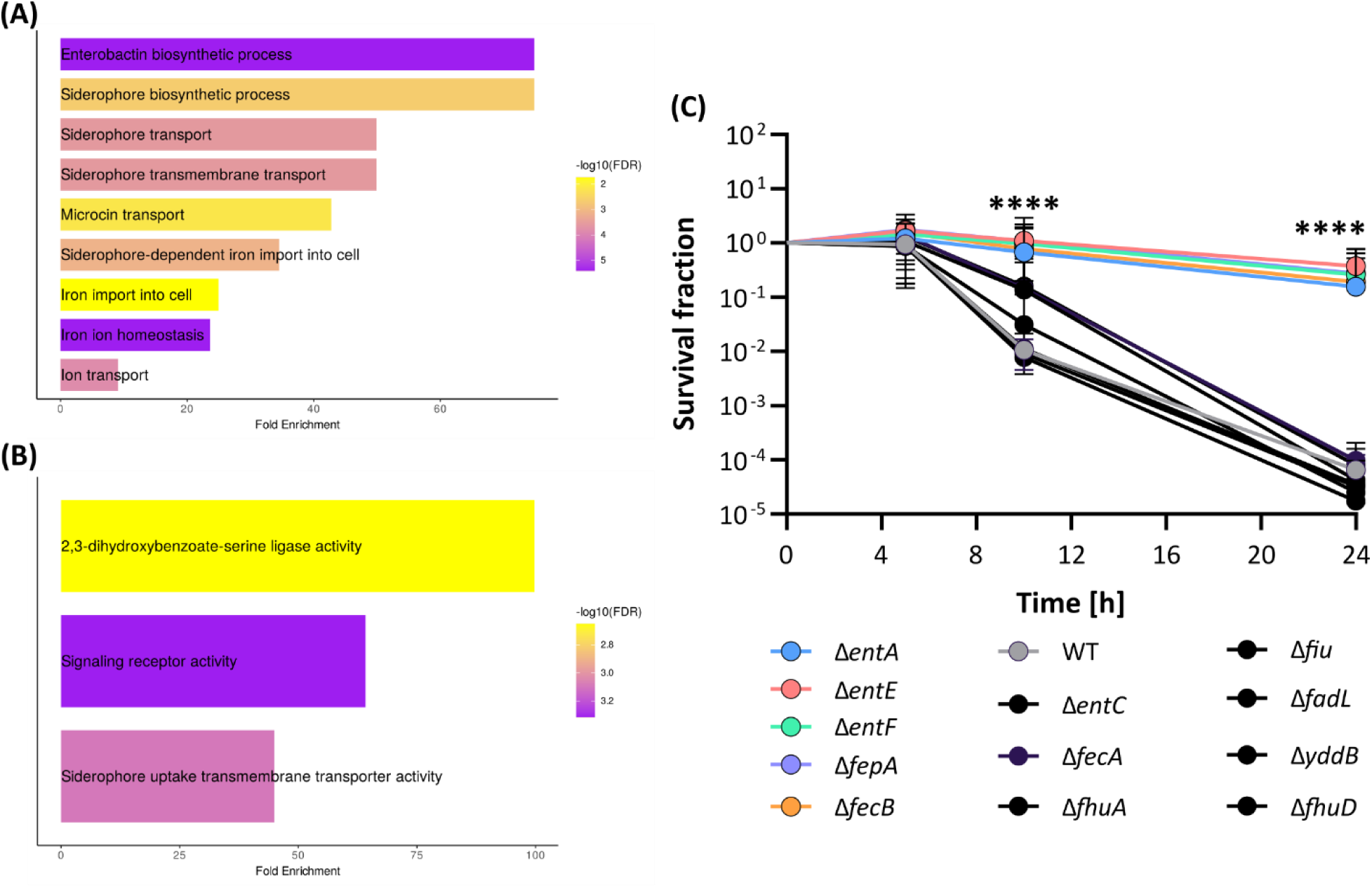
Graphical representation of proteomics-enabled enrichment analysis at the level of the **(A)** Biological Process, and **(B)** Molecular Function. Summary figures are produced from the ShinyGO application [57] (https://bioinformatics.sdstate.edu/go/). **(C)** The survival of the ion metabolism deletion mutants under 5% ethanol exposure. Colored dots represent the mean of the strains that significantly display a higher tolerance than the wild-type strain (grey). Black dots represent mutants that show the same survival as the wild-type. Error bars (n=4) represent the 95% confidence intervals and the asterisks highlight the collective *P*-value of all tolerant strains as derived from a Generalized Linear Mixed Effects Model with a Dunnett’s post-hoc test.

Next, we investigated the relevance of iron homeostasis and uptake for the ethanol tolerance phenotype. Because of the established link between ethanol toxicity and ROS stress [22,58], we hypothesized that reduced uptake of iron, as a result of impaired enterobactin (siderophore) biosynthesis (*cf.* Ent-operon) or decreased expression of dedicated iron transporters (*cf.* FecB and FepA), may explain the superior tolerance phenotype of the *envZ*\*_L116P_ tolerant strain. To assess the level of ROS stress experienced by the WT and *envZ\**_L116P_ strains under 5% ethanol stress, the fluorescence signal intensity from two distinct promoter fusion plasmids was monitored using flow cytometry [59,60]. The *soxS* promoter responds to superoxide-induced stress, triggered by 2 mM of paraquat (PQ) (Figure S17A), and the *dps* promoter senses hydrogen peroxide (Figure S17B). When cells are exposed to 5% ethanol, the alcohol significantly induces expression of the *soxS*-linked GFP signal, albeit to a lesser extent than 2 mM PQ, while induction of *dps* barely happens.

When comparing the relative induction levels in the the sensitive WT and tolerant *envZ*\*_L116P_ strain, we could not detect any difference in *soxS* or *dps* expression profile under ethanol exposure, neither under ROS stress. To revisit the link between ROS stress and ethanol tolerance, the WT and “iron metabolism attenuated”, tolerant mutants were subjected to a range of PQ and hydrogen peroxide concentrations to determine their resistance profiles towards these ROS (Figure S17C and D). This assay revealed that the more ethanol-tolerant mutants, including *envZ*\*_L116P_, could not systematically cope with ROS species better than the wild-type reference strain. Hence, despite its causal relationship with ethanol tolerance, reduced iron import does not seem to reduce the ROS stress under ethanol tolerance, which is believed to be a hallmark of ethanol tolerance, or provide any protection to ROS stress for the host.

### The tolerance-improving, L_116_P evolved allele also enhances ethanol production

Finally, we aimed to explore the potential of utilizing the *envZ*\*_L116P_-conferred ethanol tolerance to improve a strain’s ethanol production characteristics. To assess the industrial potential of the L_116_P allele, the glucose consumption and ethanol production parameters of the *envZ*\*_L116P_ mutant were tracked during an eight-day serial, fed-batch fermentation experiment and compared to those of the wild type. When analyzing the fermentation profiles, the *envZ*\*_L116P_ produced, based on glucose as substrate, ethanol faster than the ethanol-susceptible, wild-type ancestor over a 180h period (Figure 8, Figure S.17). Hence, our production results indicate that an improved tolerance phenotype, due to the L_116_P allele, also stimulates the production rate (+35%), even at ethanol concentrations below the 5% toxicity limit (the maximum EtOH concentration at the end reached 3.5-4.0 v/v%). Hence, this also suggests that bacterial production performance is even more sensitive to increasing ethanol concentrations than cellular survival.

**Figure 8:**
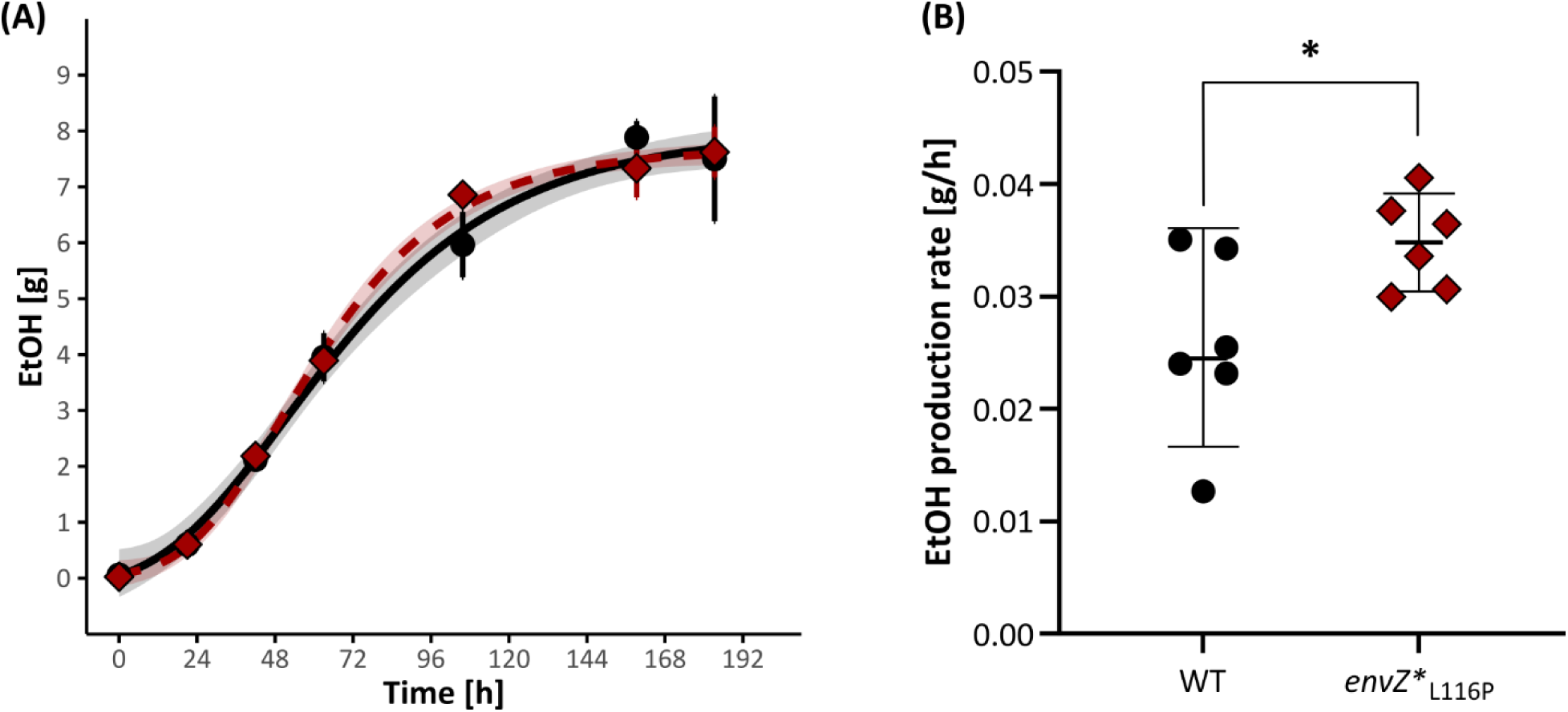
The ethanol-tolerant *envZ*\*_L116P_ displays higher ethanol production rates than the WT. **(A)** The absolute amount of ethanol (in g) in the wild type (black dots, solid line) and *envZ*\*_L116P_ strains (red diamonds, dotted line) over time. The production data are fitted using a four-parametric, sigmoidal Gompertz equation to extract the production rate value. Error bars indicate the 95% confidence interval and the ribbons surrounding the curves highlight the 95% confidence interval of the Gompertz curves as derived from a Monte-Carlo simulation. **(B)** Ethanol production rates [in g/h] in the wild type and *envZ*\*_L116P_ tolerant mutant. The production rates were statistically compared using a two-sided t-test and the level of significance is indicated as: *, *P*≤0.05, **, *P*≤0.01, ***, *P*≤0.001, ****, *P*≤0.0001.

## DISCUSSION

Exposure to high concentrations of ethanol imposes significant stress on microorganisms, leading to cellular intoxication and, ultimately, cell death. Hence, ethanol toxicity is often responsible for reducing microbial ethanol production in industrial settings, especially when the alcohol concentration accumulates over time. Therefore, improving the intrinsic tolerance of microbial production strains is an effective strategy to mitigate the adverse impact of ethanol and ensure continuous ethanol production.

Based on the mutational data from the evolutionary experiment of Swings *et al*. (2017), we identified an ethanol-tolerance improving mechanism involving a single leucine to proline substitution in EnvZ that significantly altered the balance of kinase-phosphatase activity of the enzyme. This activity is crucial for transducing osmotic stimuli towards OmpR and for engaging an osmotic response [34]. We have prioritized this system because both the sensor and the response regulator were frequently mutated in distinct, parallel-evolved populations. Moreover, the EnvZ-OmpR signal transduction pathway plays a pivotal role in the regulation of multiple membrane-associated transporters, which, according to the GO enrichment performed in this study, are required for *E. coli* to acquire ethanol tolerance. We assessed the role of the most recurrent adaptive mutation in EnvZ, L_116_P, and demonstrated that this single amino acid substitution turns *E. coli* more tolerant towards ethanol. The hypertolerant phenotype as a result of the L_116_P substitution is attributed to an improved outer membrane stability, due to an increase in total OmpC and OmpF levels, in combination with a reduction in enterobactin biosynthesis and iron uptake.

Regarding the effect of the periplasmic L_L116_P substitution, we established that this adaptive mutation, alongside T_247_R, nudges the enzymatic balance of EnvZ in favor of kinase activity. To the best of our knowledge, the majority of the mutations that affect the kinase or phosphatase reactions have been identified in the catalytic, C-terminal domain of EnvZ [38,48,61,62]. Interestingly, Yaku et *al.* (1997) were the only authors who discovered Leu residues within the periplasmic domain that upon substitution with Pro residues distort the kinase:phosphatase balance in favor of the kinase activity [63]. Hence, structural rearrangements in the periplasmic domain, as a result of amino acid substitutions, may be propagated towards the cytoplasmic domain causing perturbation of helical backbone stabilization, an altered spatial orientation of H_243_ relative to other key residues (such as D_244_), and ultimately a switch in kinase:phosphatase behavior [35].

Clearly, the observed improvement in tolerance is not merely the result of an adapted signal transduction pathway but results from the differential expression of downstream-regulated genes involved in the stress response. First, we established that porins (OmpC and OmpF) have a profound impact on the permeability and structural integrity of the outer membrane. Eliminating *ompC* reduces *E. coli*’s survival under ethanol stress, as evidenced by the appearance of morphological defects that become more pronounced with prolonged alcohol exposure. Indeed, Zhang *et al.* (2018) previously suggested a role of OmpC in ethanol tolerance in *E. coli*. Moreover, Sen *et al.* (2023) reported that alcohols cause a strong reduction in outer membrane OmpC levels, explaining the toxic effect of these chemicals on *E. coli* [64]. While we observed that eliminating *ompC* sensitizes cells, we show here that the diminished ethanol tolerance can be compensated by introducing an additional genomic copy of *ompF.* This suggests that not the porin composition, but rather the porin abundance, defines a cell’s ethanol tolerance. Moreover, the total porin content in the *envZ*\*_L116P_ evolved strain is higher than the sensitive wild-type strain, indicating that a higher OM porin content reinforces the cell envelope against ethanol exposure. Importantly, lowering the total porin content in the OM by deleting *ompC* does clearly affect the tolerance of the *envZ*\*_L116P_ mutant, but does not completely abolish it. Hence, the beneficial effect of *envZ*\*_L116P_ is not solely attributed to an increase in collective OmpC, and OmpF porin levels. Therefore, we investigated the importance of other downstream regulated genes of OmpR, resulting in the identification of TolC and LpcA as key determinants for ethanol tolerance. Interestingly, unlike *ompC*, eliminating these new targets does completely annihilate the hypertolerant phenotype of *envZ*\*_L116P_. Together with AcrAB, the TolC OM channel constitutes a tripartite efflux pump that is well known to cause antibiotic resistance by expelling antimicrobial drugs [65,66]. Previously, enhanced expression of *tolC* has been shown to confer tolerance to cyclohexane [67]. Here, we established that TolC fulfills a crucial function in a cell’s resilience to ethanol toxicity, a polar alcohol. The latter suggests that this OM channel is presumably involved in the stress response towards a broad range of chemicals and solvents, irrespectively of their hydrophobic or hydrophilic character. The second identified OmpR-regulated gene, called *lpcA*, encodes a sedoheptulose-7-phosphate isomerase that is involved in LPS synthesis. Woodruff *et al.* (2013) previously reported that overexpression of *lpcA* improves growth under ethanol stress, while Reyes *et al.* (2013) demonstrated that *lpcA* is upregulated in *n-*butanol-adapted *E. coli* cells [68,69]. Interestingly, none of the other OmpR-regulated genes influenced the tolerance level of the *envZ*\*_L116P_ mutant, although some of them were previously linked to ethanol tolerance, like the *cad* acid stress-resistant operon and the osmolyte synthesis repressor BetI [27,28]. Hence, our study demonstrates that EnvZ-OmpR-mediated adaptation to ethanol stress primarily targets OM stabilization, with the LPS synthesis enzymes LpcA and OM-embedded proteins playing a key role. Eliminating each of these membrane-stabilizing genes did affect the permeability of the OM. This observation is in agreement with previous reports highlighting the importance of interactions between LPS and outer membrane proteins in maintaining cell shape and envelope integrity under both normal physiological and stressful conditions [53,54,70]. Deleting *lpcA* results in incomplete LPS structures that may be unable to participate in cation-mediated intermolecular interactions at the OM potentially leading to (partial) collapse of this cellular component [71,72]. Hence, weakening of the OM structure, caused by a reduction of OMPs or inconsistent LPS biosynthesis, is expected to increase the susceptibility of *E. coli* to ethanol.

In addition to the OmpR-targeted deletion method, we applied an unbiased proteomics-driven approach that identified iron homeostasis as a key factor in the *envZ*\*_L116P_-associated tolerance phenotype. Reduced expression of enterobactin biosynthesis genes and iron transporters underlies the *envZ*\*_L116P_-associated tolerance. When these primary targets were knocked out in the sensitive wild-type strain, tolerance was improved. The underlying reason why reduced iron metabolism and transport are contributing to ethanol tolerance remains unclear, although it does not seem to be related to reduced ROS stress under ethanol exposure

Recently, Machas *et al.* (2021) discovered that *ompR* is also implicated in the transcriptional response of *E. coli* under exposure to styrene [73]. The latter suggests that modifying the EnvZ-OmpR signal transduction pathway, by substituting key residues in the osmosensor for instance, may provide a common strategy to achieve microbial tolerance towards a broad range of industrially-relevant hydrophilic (*e.g.* alcohols) or hydrophobic (*e.g.* styrene) solvents and chemicals. In the case of EnvZ*_L116P_, introducing this allele in an ethanol-susceptible strain improves fermentation production kinetics. This proof on concept demonstrates that tolerance engineering is relevant for microbial production, making sustainable, microbe-based manufacturing more attractive.

## MATERIALS AND METHODS

**Note**: All data analysis files are provided as R scripts (in quarto format) in the .zip folder within the same directory as the corresponding figure. The same applies for Fiji (imageJ) scripts that were used to quantitatively analyze pictures from spot assays.

### Prioritization of the EnvZ-OmpR TCS from the mutation dataset

A mutational dataset was extracted from the from the study of Swings *et al.* (2017) to identify the most predominant pathways that conferred ethanol tolerance in the ethanol-adapted *E. coli* populations [30,31]. This dataset is provided as IAMBEE_mutationSet.xlsx (Datasets folder). The entire analysis workflow is described in the R script, Fig1-IdentifyEthanolTolerancePathways.qmd. First, a mutation matrix was constructed for each of the 16 evolved *E. coli* populations (labeled from HT1-16) indicating the absence (0) or presence (1) of a certain mutation within the population. These matrices enabled summing the binary matrices to obtain the total number of mutations within a gene across all 16 evolved populations at any ethanol level, called the global pattern. Hence, if a gene was mutated multiple times, all evolved mutations within the same gene contributed to the global pattern. To retrieve the most relevant mutations (among the >2500 unique mutations), we arbitrary set the “mean occurrence” threshold at 1, meaning that a gene should carry (on average) one mutation to be retained for follow-up GO enrichment analysis. In the script, the corresponding entrez ID is linked to a specific locus or gene name since most GO enrichment tools (like GOfuncR https://www.bioconductor.org/packages/release/bioc/html/GOfuncR.html) require this input format. Linking the entrez ID to the locus or gene names was performed with the ‘bitr’ function in the clusterprofiler package (https://bioconductor.org/packages/release/bioc/html/clusterProfiler.html) and the readily available “org.EcK12.eg.db” (https://www.bioconductor.org/packages/release/data/annotation/html/org.EcK12.eg.db.html) genome-wide annotation. The script behind these operations can be found in a separate quarto file: Fig1-ExpandEntrezID.qmd. Thereafter, the enriched GO terms from the subset of genes that exceeded the mean occurrence threshold were determined using the online ShinyGO application (http://bioinformatics.sdstate.edu/go/) and, in parallel, the GOfuncR Bioconductor package. A graphical representation of the ShinyGO output is provided as supplementary figure while the output of GOfuncR is summarized in an excel-file, named GO_summary.xlsx. Both GO enrichment approaches prioritized similar GO terms, related to signaling, kinase (protein phosphorylation) activity, nucleotide binding, and predominantly restricted to the cell envelope. Therefore, we focused the analysis pipeline on two-component systems since these signal transduction pathways are commonly associated with these GO terms. To study these TCSs in more detail, we first composed a list of all known TCSs in *E. coli* based on the information provided in the KEGG database (https://www.genome.jp/kegg-bin/show_pathway?ko02020). Then, a corrected frequency score (*f*_*score*_) was assigned to each TCS-associated mutation (*m*) based on its occurrence (*n*) throughout all parallel-evolved populations and the sequencing coverage (*c*_*p*_), *i.e.* frequency, in the population-wide WGS data:

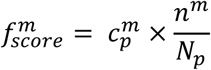

in which *N*_*p*_ represents the total number of parallel-evolved populations (16). These results are provided in frequencyTableTCS.xlsx. Next, the corrected frequency score was transformed into a global frequency score (*f̅*_*score*_) by summing the previously-calculated frequency scores of all mutations 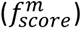 within the TCS of interest to the number of ethanol levels considered (*N*_*e*_ =7).

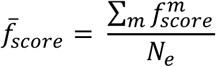

A graphical representation of the (global) frequency scores for each TCS is provided in the subfolder TCSfrequencyGraphs. Since the ShinyGO also identified membrane-related transporters to be important for ethanol tolerance, we looked into the regulons of the TCSs to prioritize those that have the highest impact on the expression of membrane-related transporters. Therefore, a list of all downstream-regulated genes for each response regulator was retrieved from the RegulonDB database (https://regulondb.ccg.unam.mx/datasets). Next, all downstream-regulated genes were GO annotated using the AnnotationDbi package to count for every TCS the number of genes encoding membrane-associated transporters (*N*_*MT*_) using the keywords: “membrane” as cellular component and “Transport” as biological process. Finally, we assigned an overall priority score (*P*_*score*_) for each TCS which is defined as the product of the global frequency scores (*f̅*_*score*_) and the number of membrane-related transporters (*N*_*MT*_):

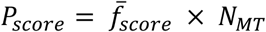

### Strains and culture conditions

The *Escherichia coli* strain BW25113 served as the reference, wild-type strain in all experiments. In case deletion mutants were included in tests, these strains were directly retrieved from the Keio library when the effect was studied in presence of wild-type *envZ* (Table M1) [74]. Otherwise, the FRT-Km^R^-FRT cassette, replacing the gene of interest, was recombined in the *envZ\**_L116P_ mutant background to examine the combined effect of the L_116_P substitution and the gene deletion on ethanol tolerance (method discussed in the next section) (Table M1). All strains were grown overnight in Lysogeny broth (LB) in an orbital shaker at 200 rpm and 37°C. As an exception, strains harboring the heat-sensitive CRISPR-FRT or pKD46 plasmids were grown at 30°C (Table M2).

**Table M.1:** List of *E. coli* strains and mutants used in this study.

| Strain | Genotype and description | Reference |
| --- | --- | --- |
| BW25113 (WT) | *rrnB3 ΔlacZ4787 hsdR514 (araBAD)567 Δ(rhaBAD)568 rph-1*, K-12 derivative | [74] |
| $\Delta envZ::Km^R$ (JW3367) | BW25113 in which the *envZ* gene is replaced by a FRT-$Km^R$-FRT resistance cassette. | [74] |
| $\Delta ompR::Km^R$ (JW3368) | Idem but the *ompR* gene is replaced by a FRT-$Km^R$-FRT resistance cassette | [74] |
| $\Delta ompC::Km^R$ (JW2203) | | [74] |
| $\Delta ompF::Km^R$ (JW0912) | | [74] |
| $\Delta tolC::Km^R$ (JW5503) | | [74] |
| $\Delta lpcA::Km^R$ (JW0212) | | [74] |
| $\Delta fhuA::Km^R$ (JW0146) | | |
| $\Delta fhuD::Km^R$ (JW0148) | | |
| $\Delta fiu::Km^R$ (JW0561) | | |
| $\Delta entF::Km^R$ (JW0578) | | |
| $\Delta entC::Km^R$ (JW0585) | | |
| $\Delta entE::Km^R$ (JW0586) | | |
| $\Delta entA::Km^R$ (JW0588) | | |
| $\Delta yddB::Km^R$ (JW1490) | | |
| $\Delta fadL::Km^R$ (JW23418) | | |
| $\Delta fecB::Km^R$ (JW4250) | | |
| $\Delta fecA::Km^R$ (JW4251) | | |
| $\Delta fepA::Km^R$ (JW5086) | | |
| *ompF' / ompF* | BW25113 strain in which the original *ompC* gene was replaced by an extra copy of *ompF* (denoted *ompF'*). | This work |
| *envZ\**<sub>L116P</sub> | BW25113 in which the Leu<sub>116</sub> residue in *envZ* was substituted for Pro. | This work |
| *envZ\**<sub>T247R</sub> | BW25113 in which the Thr<sub>247</sub> residue was substituted for Arg. | This work |
| *envZ\**<sub>F390L</sub> | BW25113 in which the Phe<sub>390</sub> residue was substituted for Leu. | This work |
| *envZ\**<sub>L116P</sub> $\Delta x::Km^R$ | Same as *envZ\**<sub>L116P</sub>, but gene *x* was replaced by a FRT-$Km^R$-FRT resistance cassette. More information about the choice of gene *x* can be found in the main text (Table 1). | This work |
| *envZ\**<sub>L116P</sub> *ompF' / ompF* | *envZ\**<sub>L116P</sub> mutant in which the original *ompC* gene was replaced by an extra copy of *ompF* (denoted *ompF'*). | This work |
| WT $\Delta lacZ::J23100$-sfGFP | Chromosomally-labeled GFP WT cells, superfolder (sf) GFP label was inserted at *lacZ* under expression of the strong, constitutive, artificial J23100 promoter. | This work |
| *envZ\**<sub>L116P</sub> $\Delta lacZ::P_{ULTRA}$-RFP | Chromosomally-labeled RFP *envZ\**<sub>L116P</sub> cells, RFP label was inserted at *lacZ* under expression of a strong, constitutive, artificial promoter derived from the ULTRA plasmid series. | This work |
| *envZ\**<sub>L116P</sub> $\Delta ompF \Delta lacZ::J23100$-sfGFP | Idem as WT $\Delta lacZ::J23100$-sfGFP but in the *envZ\**<sub>L116P</sub> $\Delta ompF$ background. | This work |
| $\Delta ompC \Delta lacZ::J23100$-sfGFP | Idem as WT $\Delta lacZ::J23100$-sfGFP but in the $\Delta ompC$ background. | This work |
| *envZ\**<sub>L116P</sub> $\Delta ompC \Delta lacZ::P_{ULTRA}$-RFP | Idem as *envZ\**<sub>L116P</sub> $\Delta lacZ::P_{ULTRA}$-RFP but in the *envZ\**<sub>L116P</sub> $\Delta ompC$ background. | This work |

**Table M.2:**
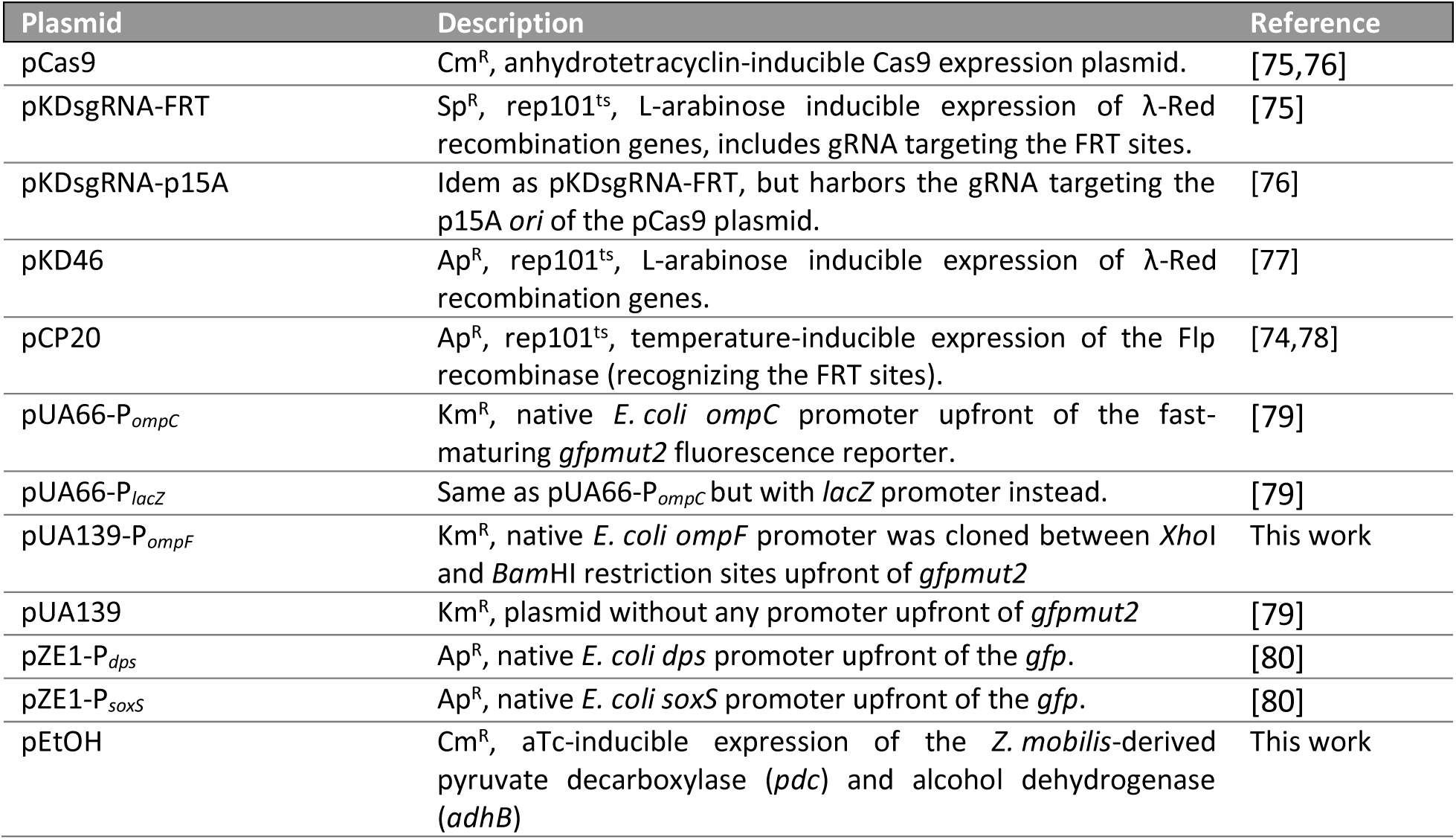
List of plasmids used in this study. Abbreviations: Cm, chloramphenicol, Km, kanamycin, Ap, ampicillin, Sp, spectinomycin, and ts, temperature-sensitive.

### Construction of deletion mutants

A recombination method based on Datsenko and Wanner [77] was applied to transfer the FRT-Km^R^-FRT cassette, that replaces the gene of interest in a specific Keio library mutant, towards the *envZ\**_L116P_ mutant. Therefore, the Km^R^-cassette was PCR amplified making sure that 200-500 bp homology overhangs were included at both sides. The Q5 PCR mix was composed according to the supplier’s protocol (New England Biolabs) and the (standard desalted) primers were ordered at Integrated DNA Technologies (Belgium) (primer sequences are listed in Table M.3). Afterward, the product was checked on a 0.7 (w/v)% agarose gel, PCR purified using the DNA Clean & Concentrator^TM^-5 (Zymo Research) and finally, quantified using the NanoDrop ND-1000. To facilitate recombination, the pKD46 plasmid that expresses the λ-Red recombination system was introduced into the *envZ\**_L116P_ mutant strain (Table M2). This plasmid was extracted from an *E. coli* TOP10 culture using the NucleoSpin® Plasmid kit (Macherey-Nagel), following the supplier’s protocol. Next, around 100 ng pKD46 was introduced into the *envZ\**_L116P_ mutant by means of chemical transformation as described by Green and Rogers (2013) [81]. When the transformation was successful, cells were incubated overnight at 30°C in the presence of 100 µg/mL ampicillin (Ap^100^) diluted in fresh LB with Ap^100^ and 0.2% L-arabinose to induce expression of λ-red genes. After 5h, cells were washed four times in 10% glycerol prior to electroporation with 200 ng PCR-amplified and purified FRT-Km^R^-FRT oligo (BioRad Pulser Xcell, 1 mm cuvette, 1.8 kV, 5 ms). Afterward, electroporated cells were recovered for 1h at 30°C and plated out on selective LB agar supplemented with 40 µg/mL kanamycin (Km^40^) and Ap^100^ for overnight incubation at 30°C. The next day, integration of the cassette at the intended locus was verified by PCR and a single correct clone was streaked on Km^40^-rich LB agar to cure the pKD46 plasmid at 42°C. After overnight incubation, loss of pKD46 was checked by spotting a couple of colonies on Ap^100^. Finally, the Km^R^-cassette was removed using the pCP20 plasmid which was introduced by electroporation (Table M2) [74]. Then, a couple of colonies were spotted on agar, supplemented with Km^40^, to verify removal of the Km^R^-cassette. Similarly to pKD46, pCP20 was cured of those clones that appear to be Km-sensitive at 42°C and loss of the pCP20 plasmid was again checked on Ap^100^.

### Reconstruction of the L_116_P, F_390_L, and T_247_R alleles in the wild-type BW25113 *E. coli* strain

The L_116_P mutation was introduced in the BW25113 strain, according to the CRISPR-FRT method as described in Swings *et al.* (2018) [75]. Briefly, the rescue oligo was retrieved from the HT11 evolved *E. coli* population, by amplifying the *envZ* locus including 500 bp up-and downstream from the mutated *evnZ* gene to facilitate recombination [30,31]. Thereafter, this rescue oligo was electroporated into the Δ*envZ*::FRT-Km^R^-FRT Keio clone harboring the pCas9 and pKDsgRNA-FRT plasmid (Table M2) [75]. In contrast to L_116_P, the F_390_L and T_247_R alleles were not readily available in our ethanol-evolved *E. coli* collection. Hence, these rescue oligos needed to be produced *de novo* using Splicing by Overlap Extension PCR (SOE-PCR) [82]. Therefore, additional primers were designed in which the overlapping 20-25 nt tails encompass the *envZ* codon to be modified. First, the fragments upstream and downstream of the desired substitution were individually amplified using the Q5 polymerase protocol (NEB). After purification, both fragments were combined in equimolar concentration (10 nM) in a 25 µL Q5 reaction mixture without additional primers. In this PCR step, 10 cycli were performed in which the PCR products were allowed to dimerize at an optimal annealing temperature, dictated by the Tm calculator tool (https://tmcalculator.neb.com/#!/main). In the second round of PCR, 25 µL Q5 mix, including the outermost primers, was supplemented to the reaction mixture and 30 cycles were run to enrich the dimerized, full-size PCR product. When the SOE PCR was completed, the product was purified using the Wizard® SV Gel and PCR Clean-Up System (Promega). This gel-purified product (*ca*. 200 ng) was again electroporated into Δ*envZ*::FRT-Km^R^-FRT strain to finish the CRISPR-FRT protocol [75]. Once these mutants were confirmed to be Km-sensitive, the *envZ* gene was amplified and sent for Sanger sequencing to confirm the presence of the desired SNP (Macrogen sequencing service). Finally, both CRISPR plasmids were either cured using an elevated growth temperature (42°C), in case of the pKDsgRNA-XXX plasmids, or using a Cas9-aided approach to remove the pCas9 plasmid according to Reisch and Prather (2015) and Swings *et al.* (2018) [75,76]

**Table M.3:**
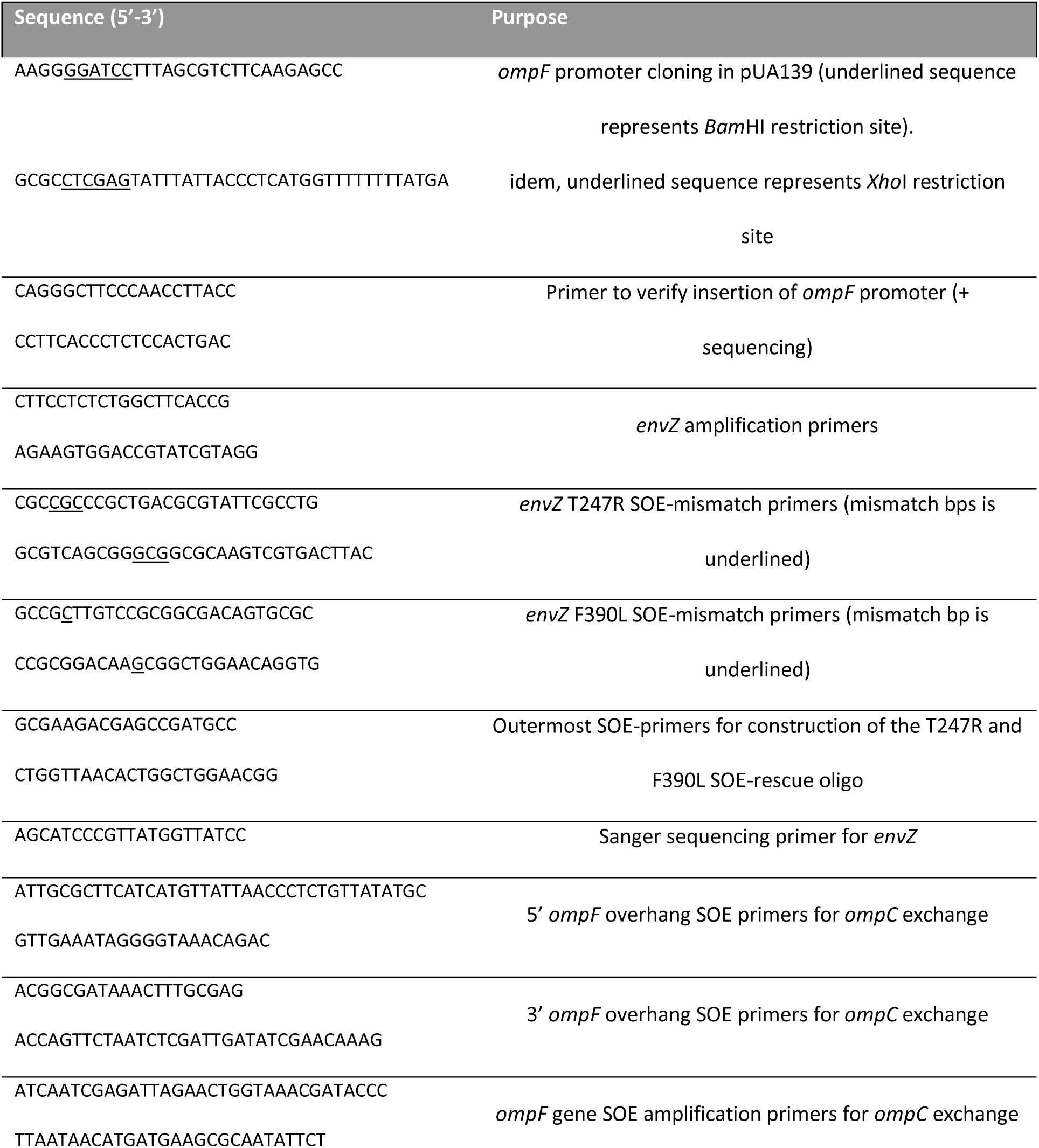
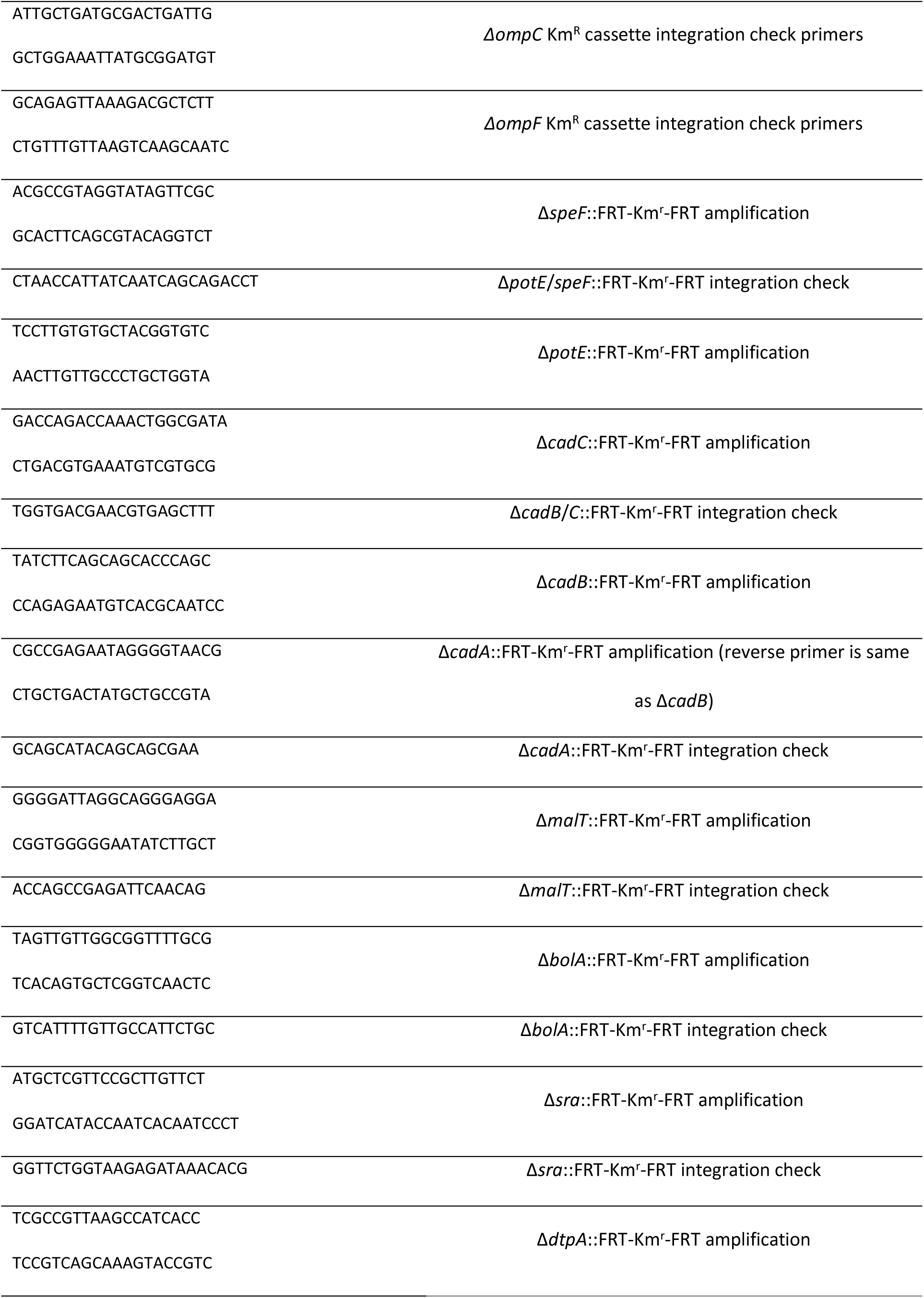
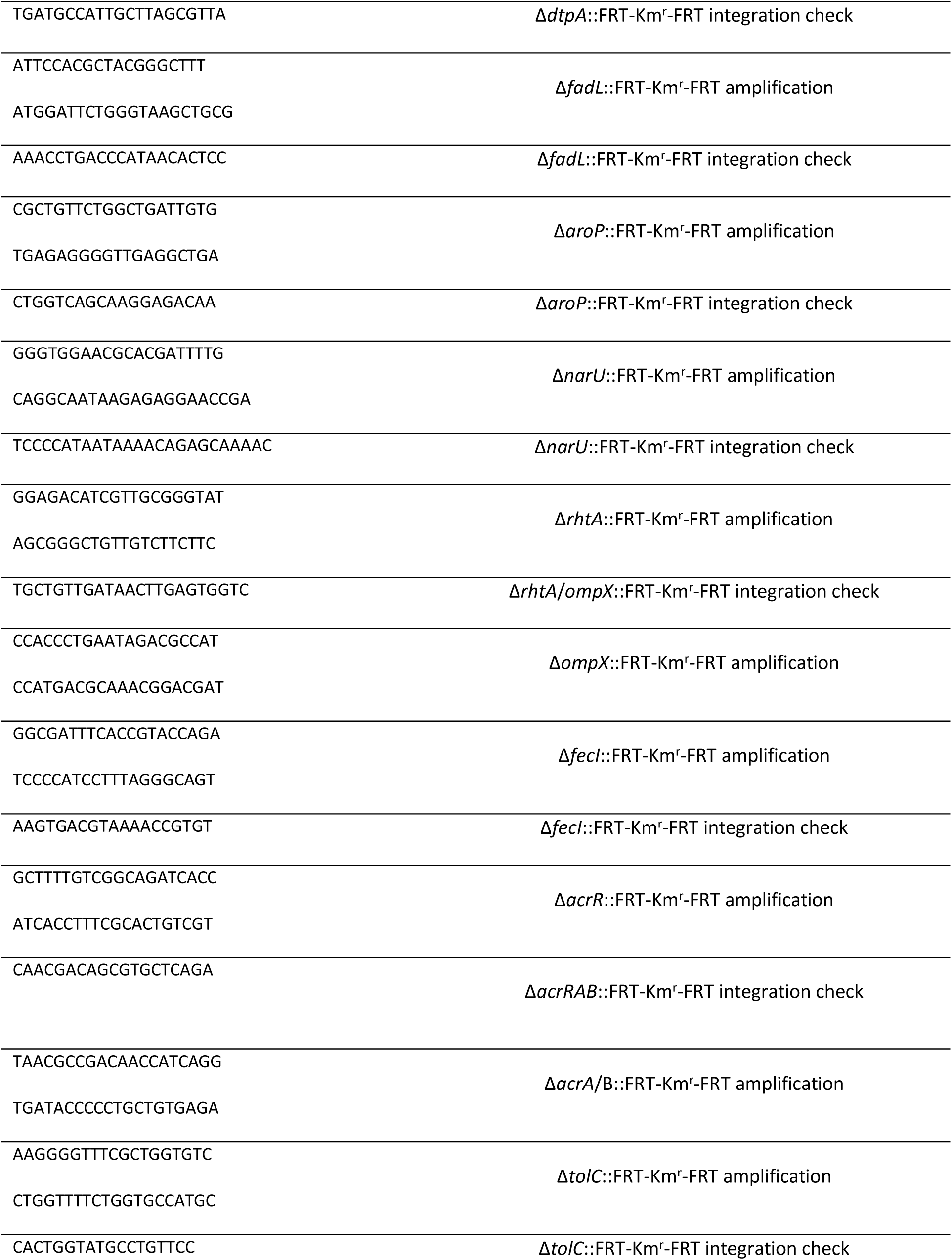
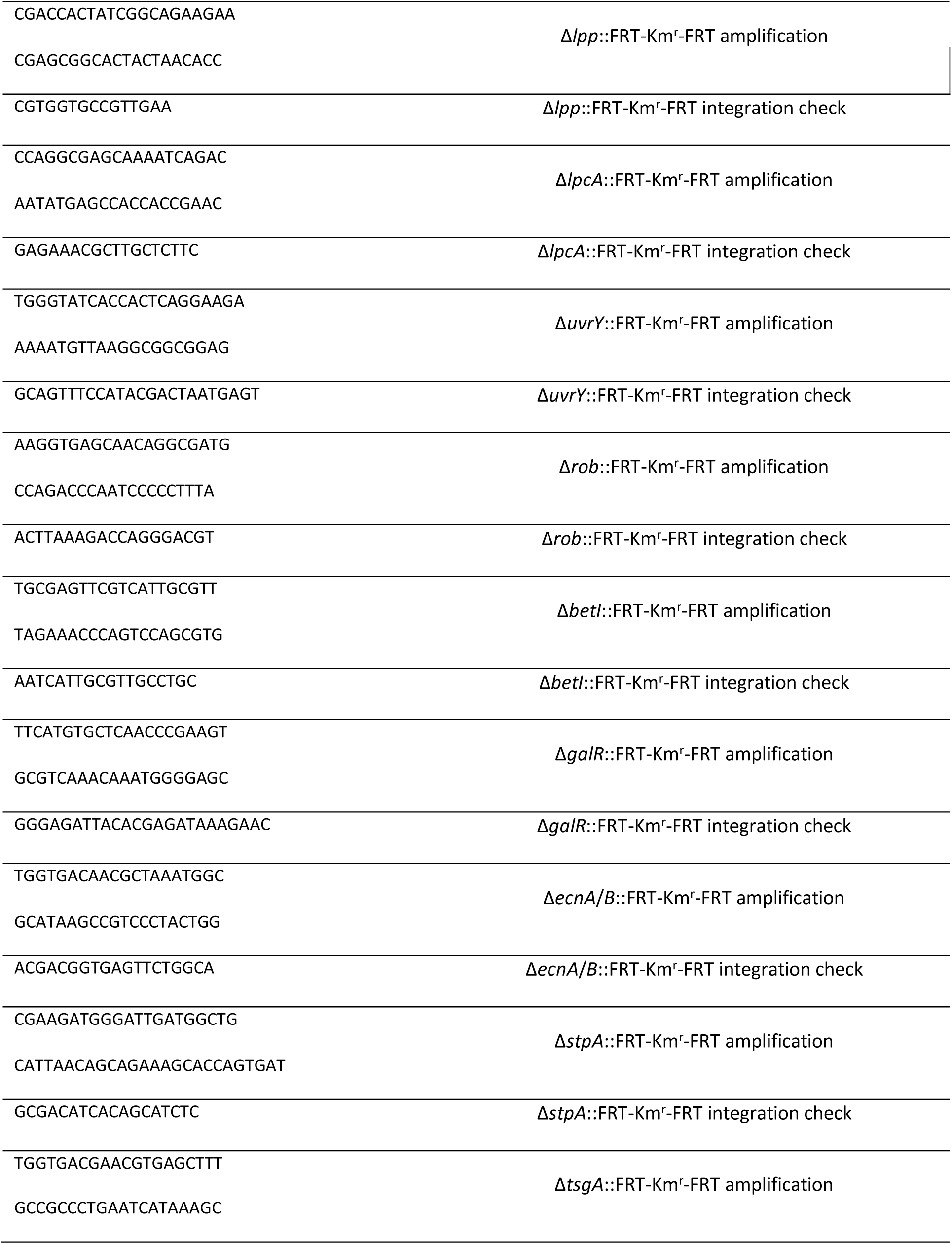
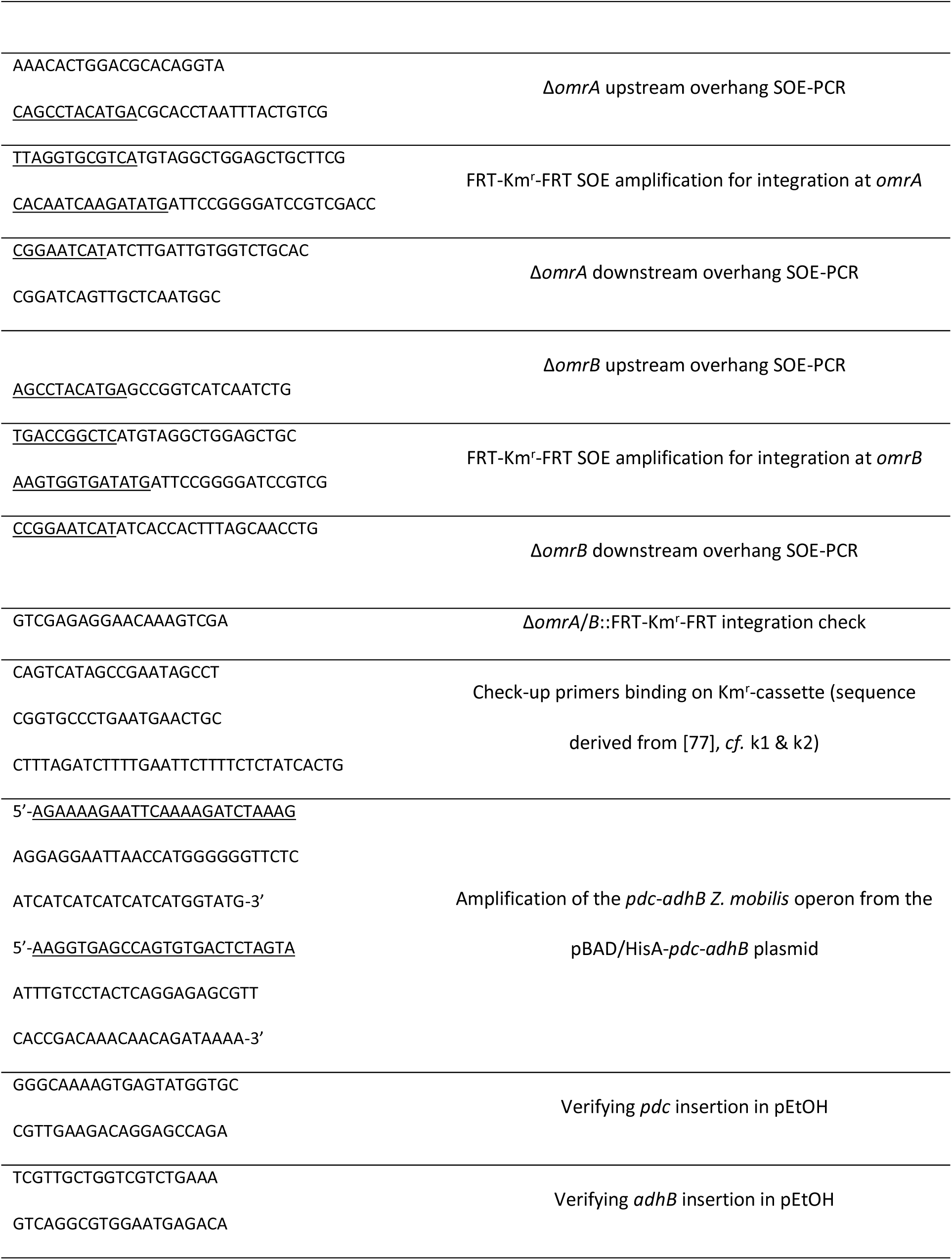
List of primers for cloning and CRISPR-FRT genome editing. Primer sequences were ordered at Integrated DNA Technologies

### Growth and survival assays in ethanol

Strains were grown overnight at 37°C in 5 mL glass tubes and the next day the optical density at 595 nm (OD_595_) was calibrated at 0.2 (ThermoFisher, Genesys 10S UV-Vis spectrophotometer) for every individual strain. Next, 2.5 mL of the calibrated cell suspension was added to each ethanol-rich flask(with screw caps to prevent ethanol evaporation) and incubated at 37°C, corresponding to a cell density of 10^6^ (which is about a 100-fold dilution). Every hour, the OD_595_ was recorded (in the first 12h interval) to retrieve the growth dynamics and survival was quantified by means of dilution plating at fixed time points (0, 5, 10, and 24h). OD-based growth curves were collected by subtracting the OD_595_ value of 5% EtOH-rich LB from a culture’s OD_595_ at any given time point. Growth parameters (including growth rate, lag time, and initial and final density) were retrieved from a spline fit, using QurvE, onto the log-transformed OD-data [41]. Finally, either the fold-increase in OD or the growth rate of the different strains were statistically compared to the wild type using a one-way ANOVA with Dunnett’s post-hoc test (using the wild-type *E. coli* or *envZ*\*_L116P_ as a reference, significance level = 0.05).

In addition, a strain’s survival fraction over 24h to 5% ethanol exposure was calculated as the ratio of the CFU count at a certain time point vs. the initial cell count. Cells were plated using the EddyJet2 spiral plater (IUL instruments) and the cell counts were automatically determined using the Flash & Grow device (IUL instruments). The survival of each mutant over the entire incubation period was compared to the reference strain using (Generalized) linear (mixed-effects) models (Time [h] x strain) with a Dunnett’s post-hoc test. Multiple models were considered and the model with the lowest Akaike Information Criterion (AIC) was selected as the most appropriate one.

### Competition tolerance assay

#### Construction of differentially labeled strains

Two fluorescent markers expressed under a strong, constitutive, artificial promoter were inserted at the *lacZ* locus in order to discriminate the competing strains. The green fluorescent sfGFP (super-folder GFP) label, was under the control of a synthetic, constitutive promoter (iGEM part BBa_K314100, designed by T. Huber and J. Spiegel (2010)) on the plasmid pSB1C3 (gift from Nicholas Coleman, University of Sydney) [83,84]. The RFP (red fluorescent protein), linked to a synthetic (Biofab) promoter, was derived from the pUltraRFP-KM plasmid [85,86]. Both promoter-fluorescent constructs were amplified from their corresponding plasmid and integrated into the *E. coli* strain of interest at the *lacZ* locus by means of SOE-PCR and homologous recombination (Table M1). Once the fluorescent constructs were successfully integrated, the Km^R^ marker was eliminated using the pCP20 plasmid as described by Cherepanov and Wackernagel (1995) (Table M2) [78].

#### Setup of the differentially-labeled, pairwise tolerance assay

For the survival-based competition assay, the fluorescently-labeled strains were individually grown overnight in LB and OD_595_-calibrated to 0.2. Then, 2.5 mL of each fluorescent strain was mixed into 50 mL 5% LB medium. For determining the survival fraction, we used the same procedure as in, but colonies were counted under the Illumatool Tunable Lighting System LT-9500 (Lightools Research) to discriminate the red and green fluorophores. For microscopy-based analysis, samples were collected every 2h (until 8h after ethanol was added to the flasks) and imaged using a Nikon Ti-E inverted microscope equipped with a Qi2 CMOS camera and 100x objective at the phase-contrast (exposure: 100 ms) and fluorescent (green: 30 ms exposure and red: 900 ms) channels. The obtained multi-channel images were processed using the MicrobeJ plugin in Fiji to extract the morphological cell parameters: cell area [µm²] and circularity [87]. In addition, each cell was characterized by fluorescence to retrieve its corresponding genotype. For statistics on morphological data, the mutant groups were first tested for normality using the Shapiro-Wilk test in Rstudio [88]. Since the datasets appeared not normally distributed, the non-parametric, Mann-Whitney test was applied at each time point [89]. Finally, the *P*-values were adjusted for multiple comparison testing using the Holm correction [90].

### Robot-assisted high throughput tolerance assay

The OmpR deletion collection was created using pKD46-mediated homologous recombination to transfer the Km-cassette from the corresponding Keio clone to the *envZ*\*_L116_ mutant strain, as described earlier. The resulting *E. coli* mutants and the *envZ*\*_L116P_ strain (Table M1) were, after calibration of cell density, treated with ethanol using the same protocol as in the standard growth and survival assay. At the initial time point (t_0_), 20 µL was sampled from each individual Erlenmeyer flask and transferred to a 96-well plate to prepare a 3-fold dilution series using the Opentrons OT-2 liquid handling robot. Then, the robot spotted 5 µL of each dilution onto a rectangular agar plate (Nunc Omnitray, Thermo Scientific). The droplets were allowed to dry before the plate was incubated overnight at 37°C. At 12h, the spot quantification procedure was repeated. The next day, the individual colonies within each spot were counted and the survival of each deletion mutant was expressed, relatively to the *envZ*\*_L116P_ reference strain that did not miss any of the genes that belong to the OmpR regulon. To identify gene deletions that were responsible for an increase/decrease in survival, we defined a linear mixed effects model (with the screened batch ID as random factor) and compared to the *envZ*\*_L116P_ reference strain using a Dunnett’s post-hoc test.Agar spot assay

### Agar spot assay

After ON incubation at 37°C, each strain was OD_595_ calibrated at 1. Then, a 10-fold dilution series (ranging from 10^0^ until 10^−6^) was prepared for each strain. Finally, 5 µL droplets were sampled from each dilution and spotted on 5 (v/v)% ethanol-rich LB agar (120 mm x 120 mm square plates, Greiner Bio-one). The agar plates were incubated at 37°C for 24h and, the next day, a picture was taken from both plates using the Canon EOS 1300D camera. This procedure was repeated four times independently. This procedure was repeated four times independently.

Afterward, the spot area and intensity were quantified using three dedicated Fiji-based scripts that are supplemented separately [91]. Briefly, a grid was superimposed onto the entire picture using the scripts ImageSplit.ijm. Using the grid as a guide, the entire picture was split into multiple sub-images, containing exactly one bacterial spot. These sub-images were stored separately for more detailed analysis in terms of spot area and pixel intensity. In case of spot area quantification, the sub-images were processed using either the SpotAnalyzer_globalTreshold.ijm script in which a particle analyzer calculated the spot area (with a surface threshold of 100 px²). In case of pixel intensity, the histogramCalculator.ijm script extracted a histogram for each sub-image in which the mean pixel intensity was included. Based on these results, the mean area and pixel intensity was determined for each bacterial spot within the entire image. Finally, a Fiji-generated image was created based on the mean spot area and intensity parameters to visually summarize the results of the four experimental repeats by means of the spotSimulator_Area&Grey_whiteBckgrd.ijm script.

### Fluorescent reporter expression assay

To estimate the bifunctional activity of the EnvZ-OmpR TCS, the promoter fusion, proposed by Zaslaver *et al.* (2006) [79], was applied. The pUA66 P*_ompC_*-*gfpmut2* and pUA66 P*_lacZ_*-*gfpmut2* constructs were directly available from the Uri Alon library (https://www.weizmann.ac.il/mcb/UriAlon/download/downloadable-data), whereas the pUA139 P*_ompF_*-*gfpmut2* was constructed ourselves (Table M2). Therefore, the *ompF* promoter (P*_ompF_*) was amplified from the BW25113 genome with primers including either the *Xho*I or *Bam*HI recognition sites. This promoter was inserted in front of the *gfpmut2* on the pUA139 plasmid, following the restriction-ligation protocol described by Zaslaver *et al.* (2006). The ligated construct was transformed into chemically competent *E. coli* TOP10 cells. Afterward, correct integration of P*_ompF_* was verified by colony PCR and sequencing. Once all three constructs were available, these three kinds of plasmids were introduced into the strains of interest and the resulting transformants were inoculated in Km-rich 5 mL LB tubes at 37°C for further testing. The next day, cells were OD_595_ calibrated at 0.2 and 2.5 mL of the cell suspension was added to each flask, filled with 50 mL 2-fold diluted LB was supplemented with Km^40^. We preferred to use 2-fold diluted growth broth to minimize background fluorescence. In case of the P*_lacZ_*-*gfpmut2* constructs, 1 mM isopropyl β-D-1-thiogalactopyranoside (IPTG) was additionally mixed into the 50 mL flasks. Every hour (up to 10h), a 150 µL sample was withdrawn from the flasks to record the fluorescent intensity (excitation: 480nm and emission: 510 nm) and OD_595_ in 96 well, F-bottom black microplates (Greiner Bio-one) using the Synergy Mx (Biotek) reader. The relative fluorescence intensity, *rF*, at time point (*t*) was expressed as:

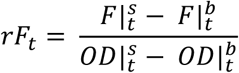

in which *OD* represents the optical density as measured at 595 nm and *s*, and *b* represent cell sample and background, respectively. The *ompC* and *ompF* expression in the wild-type and *envZ*\*_L116P_ *E. coli* strains at t=10h was quantified in relationship to the *rF* value from the *lacZ* control plasmid in each strain by dividing the *rF* linked to *ompC* or *ompF* by the *rF* associated to *lacZ*.

To measure the response of the EnvZ-OmpR TCS on osmotic stress, induced using PEG6000, or ethanol stress, the wild-type and *envZ̄*_L116P_ were cultured in 400 µL 2-fold diluted LB in a 96-deepwell plate setup (U-bottom Nunc plates, Thermo Scientific) that was shaken on an incubation platform (Heidolph Titrama× 1000, 1200 rpm, 37°C) for 3h. PEG6000 was added in concentrations of 5 or 10 (w/v)% and ethanol in 5 and 10 (v/v)%. After incubation, cells were collected by centrifugation and resuspended in 400 µL phosphate-buffered saline (PBS) (137 mM sodium chloride, 2.7 mM potassium chloride, 10 mM disodium hydrogen phosphate, and 1.8 mM monopotassium phosphate, pH 7.4). If needed, the PBS-dissolved cells were further diluted in PBS to reach a cell density of approx. 10^6^ CFU/mL and the samples were analyzed using the CytoFLEX S flow cytometer (Beckman-Coulter). Cells were discriminated from background signals using forward (FSC) and side scatter (SSC) gating and fluorescence intensity of the GFPmut2 marker was recorded at the FITC channel. The fluorescent histograms were retrieved from the Beckman-Coulter CytExpert v2.3 software and the histograms of the different conditions per strain were overlaid. For statistical comparisons, we constructed a linear model with the *ompC*:*ompF* expression ratio, expressed as 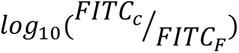, *vs.* the stress condition (w/v% PEG6000 or v/v% ethanol), in which *FITC_c_* and *FITC*_*F*_ represents the mean fluorescence intensity at the FITC channel for *ompC* or *ompF*, respectively. When the slope was significantly different from 0 (using a t-test), we considered the EnvZ osmosensor to respond to the stressor.

### Determination of OM permeability using the *N*-phenylnaphthalen-1-amine (NPN) uptake assay

Strains were incubated ON in LB tubes at 37°C in an orbital shaker (Table M1). The next day, a 12 mL bacterial stock of an OD_595_ of 0.5 was prepared in LB and three glass tubes were filled with 3 mL calibrated cell culture. To validate that OM permeabilization due to polymyxin B (PxB) treatment or ethanol could be detected, the cell cultures were incubated in presence of 0, 3, 5, 8, 10, and 15% ethanol or 0, 0.25 (1xMIC), 1.25 (5xMIC), or 2.5 µg/mL (10xMIC) PxB. Once the assay was confirmed to be suitable to quantify OM permeability, OM permeability of the strains was assayed in 0 or 5% EtOH. After 5h incubation, each of the strains was again calibrated to an OD_595_ of 0.5 in 5 mM HEPES buffer (4-(2-hydroxyethyl)-1-piperazineethanesulfonic acid, Sigma-Aldrich, pH=7.2), according to Defraine *et al.* (2018) [79]. Afterward, 150 µL of HEPES-dissolved cells were mixed with 50 µL 40 µM *N*-phenylnaphthalen-1-amine (NPN, TCI Europe, dissolved in 5 mM HEPES) and the fluorescence (λ_ex_ = 504, λ_em_ = 523 nm) and absorbance (OD_595_) were recorded using the Synergy Mx multimode reader.

The NPN uptake value of each strain under ethanol stress was defined as the NPN-derived fluorescence intensity *vs.* the optical density of the HEPES-resuspended *E. coli* culture at 595 nm. Based on these values, two statistical tests were conducted. First, a linear model was built to correlate the NPN uptake under 5% ethanol with the inherent NPN uptake values of the same strain. For *ompC*, *tolC*, and *lpcA*, the impact of deleting the target genes was corroborated with a Mixed Effects Linear Model (background x deletion) linked to a Dunnett’s post-hoc test with either the WT or *envZ*\*_L116P_ strain as reference. Finally, the relation between survival and NPN uptake was demonstrated using a Spearman correlation test.

### Whole-genome proteomics

#### Sample preparation

Overnight bacterial cultures were diluted 100-fold and inoculated in 50 mL LB for *ca.* 3.5h until anOD_595_ of 0.8 was reached. For each strain, 15 mL culture was collected and washed three times in PBS. Then, the cell pellets were frozen in liquid nitrogen for shipment to the VIB proteomics core (Ghent, Belgium). In the VIB proteomics core facility, the ell pellets were homogenized in 100 µl lysis buffer containing 5% sodium dodecyl sulfate (SDS) and 50 mM triethylammonium bicarbonate (TEAB), pH 8.5. Next, the resulting lysate was transferred to a 96-well PIXUL plate and sonicated with a PIXUL Multisample sonicator (Active Motif) for 5 minutes with default settings (Pulse 50 cycles, PRF 1 kHz, Burst Rate 20 Hz). After centrifugation of the samples for 15 minutes at maximum speed at room temperature (RT) to remove insoluble components, the protein concentration was measured by bicinchoninic acid (BCA) assay (Thermo Scientific) and from each sample 100 µg of protein was isolated to continue the protocol. Proteins were reduced and alkylated by addition of 10 mM Tris(2-carboxyethyl)phosphine hydrochloride and 40 mM chloroacetamide and incubation for 10 minutes at 95°C in the dark. Phosphoric acid was added to a final concentration of 1.2% and, subsequently, samples were diluted 7-fold with binding buffer containing 90% methanol in 100 mM TEAB, pH 7.55. The samples were loaded on the 96-well S-Trap™ plate (Protifi), placed on top of a deepwell plate, and centrifuged for 2 min at 1,500 x g at RT. After protein binding, the S-trap™ plate was washed three times by adding 200 µl binding buffer and centrifugation for 2 min at 1,500 x g at RT. A new deepwell receiver plate was placed below the 96-well S-Trap™ plate and 50 mM TEAB containing trypsin (1/100, w/w) was added for digestion overnight at 37°C. Using centrifugation for 2 min at 1,500 x g, peptides were eluted in three times, first with 80 µL 50 mM TEAB, then with 80 µL 0.2% formic acid (FA) in water and finally with 80 µL 0.2% FA in water/acetonitrile (ACN) (50/50, v/v). Eluted peptides were dried completely by vacuum centrifugation. Samples were dissolved in 100 µL 0.1% TFA in water/ACN (98:2, v/v) and desalted on a reversed phase (RP) C18 OMIX tip (Agilent). The tip was first washed 3 times with 100 µL pre-wash buffer (0.1% TFA in water/ACN (20:80, v/v)) and pre-equilibrated 5 times with 100 µL wash buffer (0.1% TFA in water) before the sample was loaded on the tip. After peptide binding, the tip was washed 3 times with 100 µL of wash buffer and peptides were eluted twice with 100 µL elution buffer (0.1% TFA in water/ACN (40:60, v/v)). The combined elutions were transferred to HPLC inserts and dried in a vacuum concentrator.

#### LC-MS/MS analysis

Peptides were re-dissolved in 20 µL loading solvent A (0.1% trifluoroacetic acid in water/acetonitrile (ACN) (99.5:0.5, v/v)) of which 2 µL of the sample was injected for LC-MS/MS analysis an on an Ultimate 3000 Pro Flow nanoLC system in-line connected to a Q Exactive HF mass spectrometer (Thermo). Trapping was performed at 20 μL/min for 2 min in loading solvent A on a 5 mm trapping column (Thermo scientific, 300 μm internal diameter (I.D.), 5 μm beads). The peptides were separated on a 250 mm Aurora Ultimate, 1.7 µm C18, 75 µm inner diameter (Ionopticks) kept at a constant temperature of 45°C. Peptides were eluted by a non-linear gradient starting at 0.5% MS solvent B reaching 26% MS solvent B (0.1% FA in acetonitrile) in 75 min, 44% MS solvent B in 95 min, 56% MS solvent B in 100 minutes followed by a 5-minute wash at 56% MS solvent B and re-equilibration with MS solvent A (0.1% FA in water). The mass spectrometer was operated in data-independent mode, automatically switching between MS and MS/MS acquisition. Full-scan MS spectra ranging from 375-1500 m/z with a target value of 5E6, a maximum fill time of 50 ms and a resolution at of 60,000 were followed by 30 quadrupole isolations with a precursor isolation width of 10 m/z for HCD fragmentation at an NCE of 30% after filling the trap at a target value of 3E6 for maximum injection time of 45 ms. MS2 spectra were acquired at a resolution of 15,000 at 200 m/z in the Orbitrap analyser without multiplexing. The isolation intervals ranging from 400– 900 m/z were created with the Skyline software tool. The polydimethylcyclosiloxane background ion at 445.120028 Da was used for internal calibration (lock mass) and QCloud [92,93] was used to control instrument longitudinal performance during the project.

#### - Data analysis

LC-MS/MS runs of all samples were searched together using the DiaNN algorithm (version 1.8.1), library free. Spectra were searched against the *Escherichia coli* protein sequences in the Uniprot database (database release version of January 2024), containing 4,403 sequences (www.uniprot.org), supplemented with the universal protein contaminant database (database release version of 2023_02), containing 381 sequences [94]. Enzyme specificity was set as C-terminal to arginine and lysine, also allowing cleavage at proline bonds with a maximum of two missed cleavages. Variable modifications were set to oxidation of methionine residues and acetylation of protein N-termini while carbamidomethylation of the cysteine residues was set as fixed modifications. Mainly default settings were used, except for the addition of a 400-900 m/z precursor mass range filter and MS1 and MS2 mass tolerance was set to 15 and 20 ppm respectively. Further data analysis of the shotgun results was performed with an in-house script in the R programming language, version 4.2.2. Protein expression matrices were prepared as follows: the DIA-NN main report output table was filtered at a precursor and protein library q-value cut-off of 1 % and only proteins identified by at least one proteotypic peptide were retained. After pivoting into a wide format, iBAQ intensity columns were then added to the matrix using the DIAgui’s R package (https://rdrr.io/github/mgerault/DIAgui/man/DIAgui.html) get_IBAQ function. PG.Max LFQ intensities were log2 transformed and replicate samples were grouped. Proteins with less than three valid values in at least one group were removed and missing values were imputed from a normal distribution centered around the detection limit (package DEP, [95]) leading to a list of 2,728 quantified proteins in the experiment, used for further data analysis. To compare protein abundance between pairs of sample groups (WT vs envZ; envZ vs envZ-ompF_ompF; WT vs envZ-ompF_ompF sample groups), statistical testing for differences between two group means was performed, using the package limma [96]. Statistical significance for differential regulation was set to a false discovery rate (FDR) of <0.05 and fold change of >4- or <0.25-fold (|log2FC| = 2). Results are provided in Fig8 - proteomics_data.xlsx.

Significantly differentially expressed proteins in the envZ *vs*. the WT group were retained for further GO enrichment analysis. Significantly enriched GO terms were identified using three similar approaches based on the Bioconductor packages, GOfuncR and clusterProfiler, and the online-accessible ShinyGO application. All three approaches prioritized the same key GO terms. Next, each protein *x* within a sample was normalized using the available log_2_.PG.MaxLFQ (log_2_-transformed maximal label-free quantification) values as:

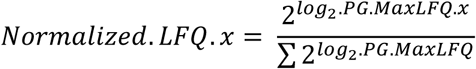

This approach enables comparing the relative protein abundances, represented by their *Normalized*. *LFQ* values, between the WT, *envZ*\*_L116P_, and *envZ\**_L116P_ *ompF’*/*ompF* strains using a one-way ANOVA with Tukey’s post-hoc test.

### Quantification of the ROS response under ethanol, paraquat, and hydrogen peroxide stress

Before the assay, the ROS reporter plasmids (pZE1-P*_dps_*and pZE1-P*_soxS_*) were introduced in the WT and *envZ*\*_L116P_ strains using heat shock transformation. Strains were cultured overnight in Ap100-rich LB medium and calibrated to OD_595_ of 0.2. In case of ethanol, the strains were diluted in fresh 5% ethanol-rich LB, just like in the growth and survival assay. When paraquat (PQ) or hydrogen peroxide (H_2_O_2_) were used, the same relative volume of calibrated cells was added to 5 mL LB, containing 2 mM PQ or H_2_O_2_. In the non-challenged condition, ethanol or the ROS molecules were replaced with demi-water. The fluorescence intensities of the promoter-linked GFP reporter were recorded using the CytoFLEX S flow cytometer at the start (t=0h) and after 1h of exposure to ethanol, PQ, or H_2_O_2_. Then, we expressed the *log*_2_(induction) for each reporter plasmid construct and condition as:

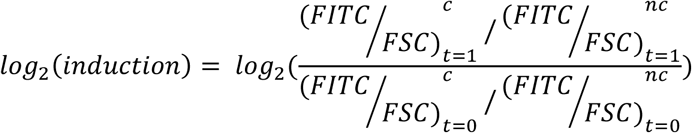

in which *FITC*, represents the FITC channel, *FSC*, the forward scatter, *c*, the challenged condition, and *nc*, the non-challenged condition. The induction levels were pairwise compared using *t-*test statistics.

### EC_50_ determination of strains for paraquat and hydrogen peroxide

After overnight incubation, strains were calibrated to OD_595_ 0.2 and 400-fold diluted in fresh LB. The cells were challenged to a 2-fold dilution gradient of H_2_O_2_ (ranging from 8 mM to 0.125 mM) or PQ (ranging from 10 to 0.156 mM). In each assay, non-treated controls were included as well. Each well of the 96-well plate was covered with silicone oil (Fisher scientific) to prevent evaporation and the plate was inserted in a Cerillo Stratus Microplate Reader. The OD_595_ was periodically recorded every 15 min for 48h at 37°C. The OD-data were processed using spline models in QurvE [41] and the growth rate was extracted. This growth parameter served as input for dose-response curve analysis (using the build-in *drc* functionality in QurvE) from which the EC_50_ values could be retrieved. The EC_50_-values of the different strains were statistically compared to the wild type using a generalized mixed effects linear model with Dunnett’s post-hoc test.

### Fermentation setup for ethanol production

To promote ethanol production in *E. coli*, the *Z. mobilis* pet operon genes (including *pdc_Zm_* and *adhB_Zm_*) from the previously constructed pBAD/HisA-*pdc_Zm_*-*adhB_Zm_*plasmid (Toon Swings, unpublished results) were transferred to the pCas9 plasmid, thereby replacing the original Cas9 nuclease. Therefore, the pCas9 plasmid was PCR linearized and the *pdc_Zm_* - *adhB_Zm_* operon was amplified. Both PCR products were equimolarly mixed for Gibson assembly using the NEBuilder HiFi DNA Assembly Master Mix. The resulting pEtOH construct was transformed into chemocompetent *E. coli* TOP10 cells, plasmid purified, and sent for sequencing (Macrogen). Finally, the pEtOH plasmid was introduced into the wild-type and *envZ*\*_L116P_ strains using heat-shock transformation.

Before fermentation, the pEtOH-carrying strains were inoculated in LB tubes, enriched with Cm^30^, and incubated overnight at 30°C. The next day, the ON cultures were diluted 100-fold into flaks with fresh Cm^30^-enriched LB medium. After 2h of incubation, the *Z. mobilis* ethanol-producing enzymes were induced with 100 ng/mL aTc for *ca.* 20h. Thereafter, cells were collected by centrifugation, the supernatant was discarded and the pellet was resuspended in 50 mL fermentation broth (Table M.4, step I). Before the flasks were sealed off with an airlock system (Brouwland), a 1 mL sample was taken for HPLC analysis. During the next days, the addition of fermentation broth and sampling was repeated 3-times with 24h in between each cycle. After the final step, the fermentation process was allowed to continue at 30°C for a total period of 185h. This fermentation setup was repeated six times for each strain.

**Tabel M.4:** Composition of the fermentation broth. Phospate buffer consisted of pH 7-buffered 93.5 mM dipotassium hydrogenphosphate and 6.5 mM monopotassium phosphate. The glucose feedstock was prepared by mixing 214 g glucose.monohydrate in 1L phosphate buffer, followed by filter-sterilization.

| Step | LB (10x) | Phosphate buffer | Glucose feedstock |
| --- | --- | --- | --- |
| I | 5.0 mL | 38.6 mL | 6.4 mL |
| II | 5.0 mL | 25.7 mL | 19.3 mL |
| III | 5.0 mL | 12.9 mL | 32.1 mL |
| IV | 5.0 mL | 0.0 mL | 45.0 mL |

The ethanol and glucose concentration was measured using an Agilent HPLC 1200 series, equipped with a Bio-Rad Aminex HPX-87H column (temperature: 55 °C) and a refractive index detector (RID, temperature 35°C). For the analysis, a degassed 1 mM sulfuric (VWR) acid mobile phase with a flow rate of 0.6 mL/min was used. For each sample, the glucose and ethanol concentrations were inferred from calibration curves.

Finally, the cumulative amount of ethanol (in g), produced by the wild-type and *envZ*\*_L116P_ mutant, was fitted with the four-parametric sigmoidal Gompertz equation (NLS.G4) to determine the production rate [in h^−1^]. This fermentation parameter was then used to statistically compare the ethanol productivity of the *envZ*\*_L116P_ *vs.* the wild-type strain using a two-sided pairwise *t*-test.

## ACKNOWLEDGEMENTS

The authors would like to thank the VIB proteomics core for performing the LC-MS/MS analysis of the *E. coli* proteome and conducting the related data analysis. Furthermore, the authors acknowledge Dr. Philip Ruelens (Centre of Microbial and Plant Genetics, KU Leuven, Leuven, Belgium; and Center for Microbiology, VIB-KU Leuven, Leuven, Belgium) for his valuable advice on the statistical processing of the survival data.

